# Novel antimicrobial activity of photoactive compounds originally synthesized for industrial applications

**DOI:** 10.1101/2025.01.21.634129

**Authors:** José Manuel Ezquerra-Aznárez, Raquel Alonso-Román, Ainhoa Lucía, Raquel Andreu, Santiago Franco, José A. Aínsa, Santiago Ramón-García

## Abstract

The emergence of antimicrobial resistance threatens advances achieved by medicine in the last century. This situation has been aggravated by the suboptimal outcome of screening campaigns to provide novel antibiotics. In this study, we took an alternative approach to discover new chemical scaffolds with antimicrobial activity. We screened a collection of photoactive compounds originally synthesized for industrial purposes and found that 4*H*-pyran-4-ylidenes are active against Gram-positive bacteria. Compounds belonging to this family displayed dose-dependent bactericidal activity against wild-type *Staphylococcus aureus* and Methicillin Resistant *S. aureus* (MRSA). Importantly, they were not cytotoxic to hepatic cell lines at the concentrations at which antimicrobial activity was observed. Resistance to 4*H*-pyran-4-ylidenes in *S. aureus* was achieved via mutations in the *rny* locus, likely via changes in mRNA levels of genes critical for their activity. Overall, we demonstrate that chemical libraries not originally intended for drug discovery can be a fruitful source of chemical diversity for the development of novel antimicrobials.

## Introduction

The discovery of antimicrobials was one of the milestones of humankind in the 20^th^ century. Their introduction brought unprecedented success in the treatment of infectious diseases and set up the development of other areas of medicine, such as invasive surgery and cancer chemotherapy. The current antimicrobial resistance (AMR) crisis threatens all these advances^1^. Nowadays, AMR kills an estimated 700,000 people every year and it is projected that, at the current pace, it will be accountable for 10 million deaths and an estimated economic loss of $100 trillion per year by 2050 ^2^. Therefore, there is an urgent need to discover and develop new antimicrobials.

Most of the currently available antimicrobial families were discovered in the mid-20^th^ century, during a period known as the Golden Age of Antimicrobial Discovery. During that time, new antimicrobials were identified by screening soil-dwelling microorganisms for activity against pathogenic bacteria. This platform eventually collapsed when the active compounds found were already known antibiotics or molecules with significant toxicity issues^3,4^. The paradigm changed again at the end of the 20^th^ century, when the first bacterial genomes became available^5^. Genomic information allowed the identification of essential targets conserved across different bacterial species and target-based approaches were developed to find active molecules against them. However, no novel classes of antimicrobials have been identified using such approaches^6,7^.

Several factors can explain the failure of molecules identified through target-based approaches for inhibiting microbial growth. First, the bacterial cell wall is a formidable barrier that protects bacteria from environmental threats, including antimicrobials^8^. Because of this, molecules that are good inhibitors of cytosolic targets might be not active against whole bacteria, thus limiting the success of target-based approaches in bacteria. An additional problem is the design of chemical libraries, which has been strongly influenced by Lipinski’s rules in the past decades^9^. Antimicrobials as a group, however, represent an exception to Lipinski’s rules, displaying higher molecular weights and polarity. Thus, chemical libraries used to run high throughput screening (HTS) campaigns (either as target-based approaches or as whole cell screenings) for antimicrobial discovery were biased and did not fully reflect the chemical diversity of current antimicrobial families^10–12^.

In this study, we sought unexplored chemical diversity and screened a collection of organic dyes synthesized for solar energy production purposes^13,14^ against a panel of pathogenic bacteria from the World Health Organization priority list^15^. We found that 4*H*-pyran-4-ylidene derivatives were active against Gram-positive bacteria. Compounds belonging to this family were bactericidal to *Staphylococcus aureus*, including an MRSA strain, and non-toxic to mammalian cells nor mutagenic to a tryptophan-auxotrophic *Escherichia coli* strain.

## Results

### Identification of antimicrobial compounds in a panel of Photoactive Molecular Materials (PMM)

We screened a panel of 21 compounds representing the chemical diversity from the PMM compound collection for antimicrobial activity against *Mycobacterium tuberculosis* H37Rv. One of them, a 4*H*-pyran-4-ylidene (compound 01, **Table S1**), was active at 50 µM. We then tested a panel of 39 compounds belonging to the 4*H*-pyran4-ylidene family (**Table S2**, compounds 01-39) against a set of Gram-positive, Gram-negative, and non-tuberculous mycobacterial strains (**Table S3**). Of them, thirteen were active against at least one bacterial species at 50 µM (**Table S2, Table S4).**

We subsequently validated the activity of the 13 compounds identified as active against at least one bacterial species in a dose-response assay aimed at determining their Minimal Inhibitory Concentration (MIC) and Minimal Bactericidal Concentration (MBC) (**Table 1**). Four compounds—04, 13, 15, and 36— were inactive against all Gram-positive strains, thus considered false positives from the single-shot assay; these were discarded for further assays. The nine remaining compounds were classified into two different groups: compounds with broad-spectrum activity against several Gram-positive species (02, 11, 18, 19, and 27) (**Fig. 1**), and compounds whose activity was restricted to just one out of the six bacterial species tested (05, 07, 13, and 39). MBC matched MIC values in most cases, strongly suggesting that 4*H*-pyran-4-ylidene derivatives are bactericidal to Gram-positive bacteria.

**Fig. 1.**
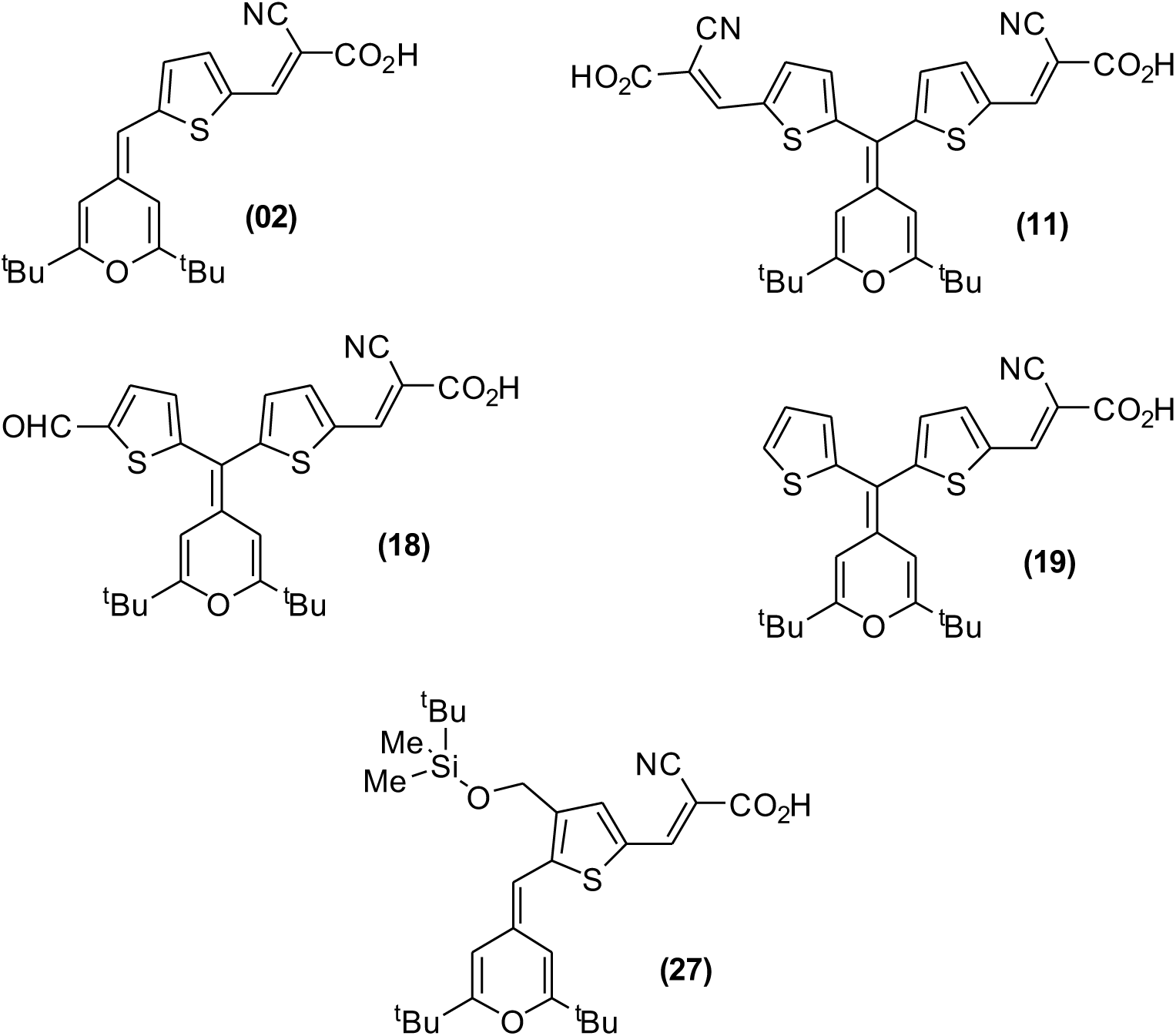
Chemical structures of 4*H*-pyran-4-ylidene derivatives with broad-spectrum activity against Gram-positive species.

**Table 1.**
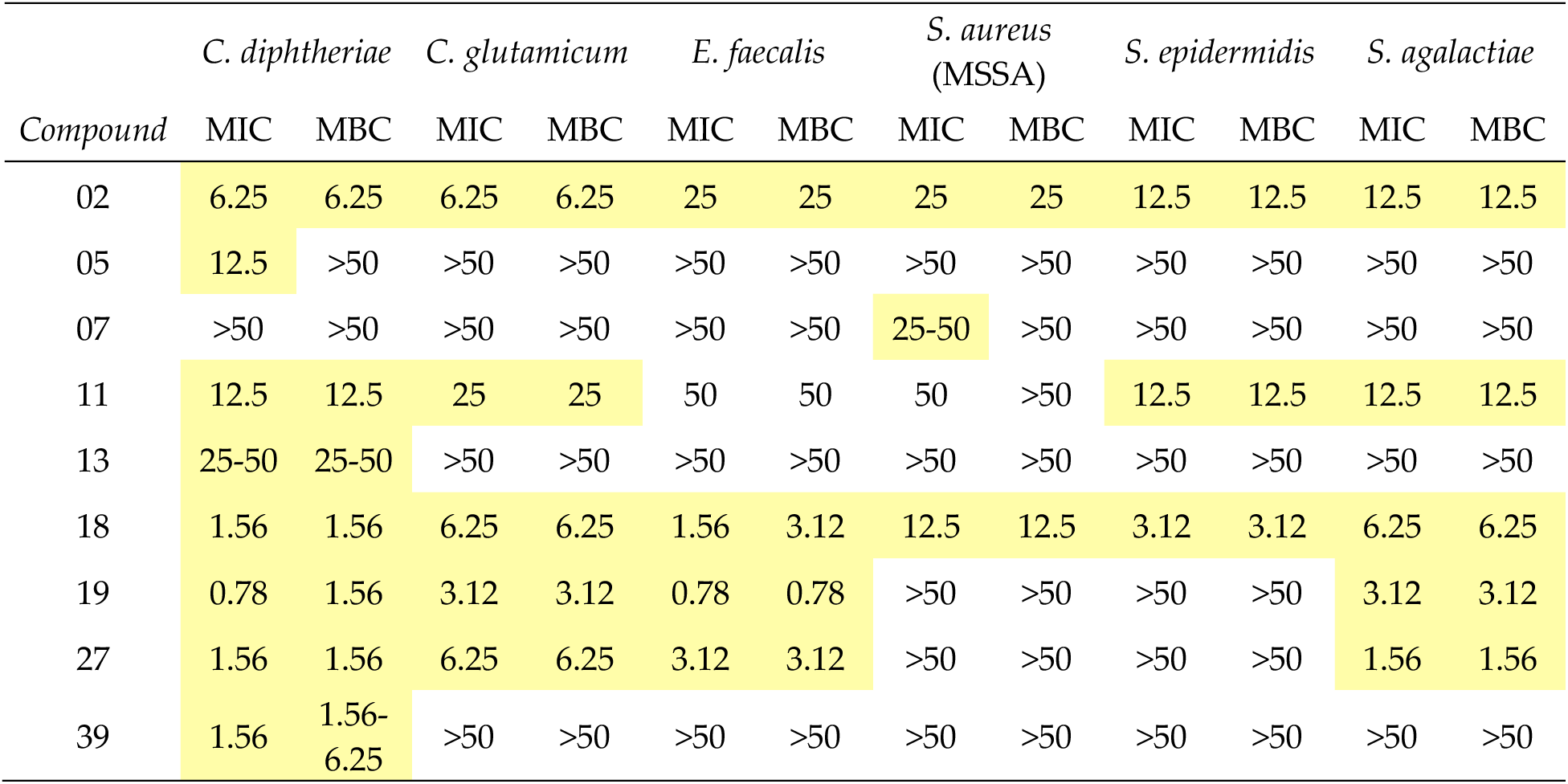
Minimal inhibitory and bactericidal concentrations for the nine active compounds tested in dose-response assays. Concentrations are indicated in µM. MIC/MBC values below 50 µM are indicated in yellow.

### Compound 18 is bactericidal to *S. aureus*

In order to confirm the potential bactericidal activity of 4*H*-pyran-4-ylidene derivatives suggested by the dose-response MIC/MBC assays, we performed time-kill kinetics of Compound 18 against two *S. aureus* strains: ATCC 29213 (methicillin-susceptible, MSSA, MIC = 25 µM) and ATCC 43300 (methicillin-resistant, MRSA, MIC = 12.5 µM). Indeed, Compound 18 exhibited bactericidal activity against both strains, achieving maximal killing at the 6-hour timepoint for concentrations equal to or greater than the MIC (**Fig. 2**). Subinhibitory concentrations were also able to reduce the bacterial population at early timepoints, but growth was restarted after 1 hour. Compound 18 seemed to be slightly more potent against the MRSA strain, which was unable to resume growth in the presence of the maximum concentration tested, 50 µM.

**Fig. 2.**
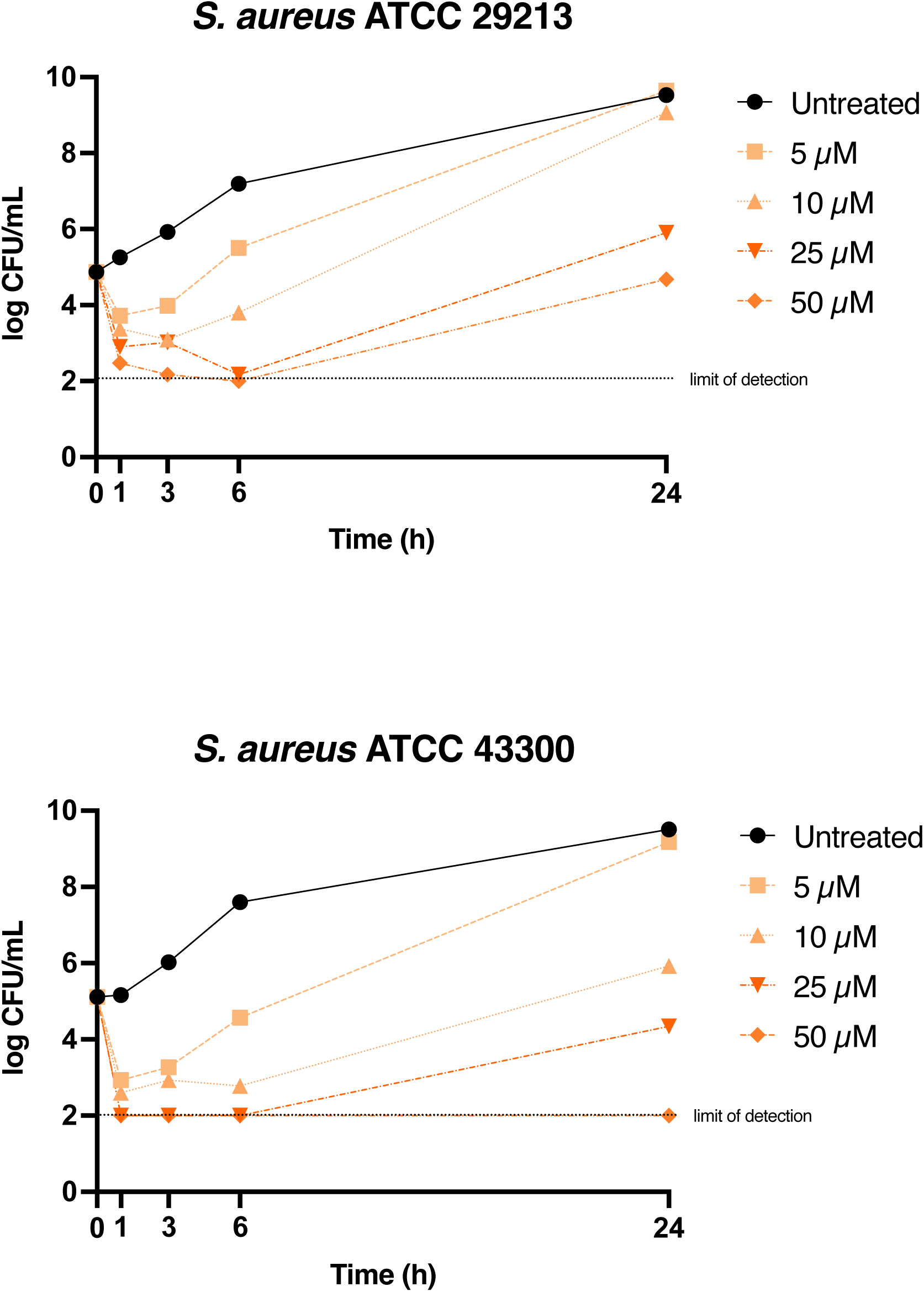
Time-kill kinetics of Compound 18 against *Staphylococcus aureus*. Two strains were used: *S. aureus* ATCC 29213 (methicillin-susceptible, MIC = 25 µM) and *S. aureus* ATCC 43300 (methicillin-resistant, MIC = 12.5 µM).

In order to evaluate whether the outgrowth was caused by the emergence of stable genetic resistance or transient phenotypic adaptions, we subcultured the populations exposed to 25 µM of Compound 18 and subsequently tested their susceptibility to it. MIC determinations showed that the methicillin-susceptible *S. aureus* ATCC 29213 population contained mutants with decreased susceptibility to compound 18, as the population MIC increased to 100 µM. The population derived from the MRSA strain, on the other hand, remained susceptible to Compound 18.

### Mutations in the *rny* locus confer *S. aureus* resistance to Compound 18

We were able to isolate *S. aureus* MSSA mutants resistant to Compound 18 from two independent experiments. First, cells treated with 25 µM (1x MIC) of Compound 18 for 24 hours were allowed to recover in fresh medium and, subsequently, exposed to 100 µM (4x MIC) of Compound 18 for a further 24 hours. Then, the culture was inoculated onto compound-free plates and individual clones selected for MIC assays. Second, we isolated mutants directly by seeding a culture onto plates containing 100 µM of Compound 18; this approach demonstrated a mutant frequency of 7·10^-7^ of Compound 18. In both cases, we were able to isolate mutants with MIC increases of 4-fold or higher (i.e. 100 µM). We then validated their resistant phenotype using a drop-dilution assay to confirm that the mutants were able to grow in the presence of up to 200 µM of Compound 18 (**Fig. 3** and **Fig. S1**).

**Fig. 3.**
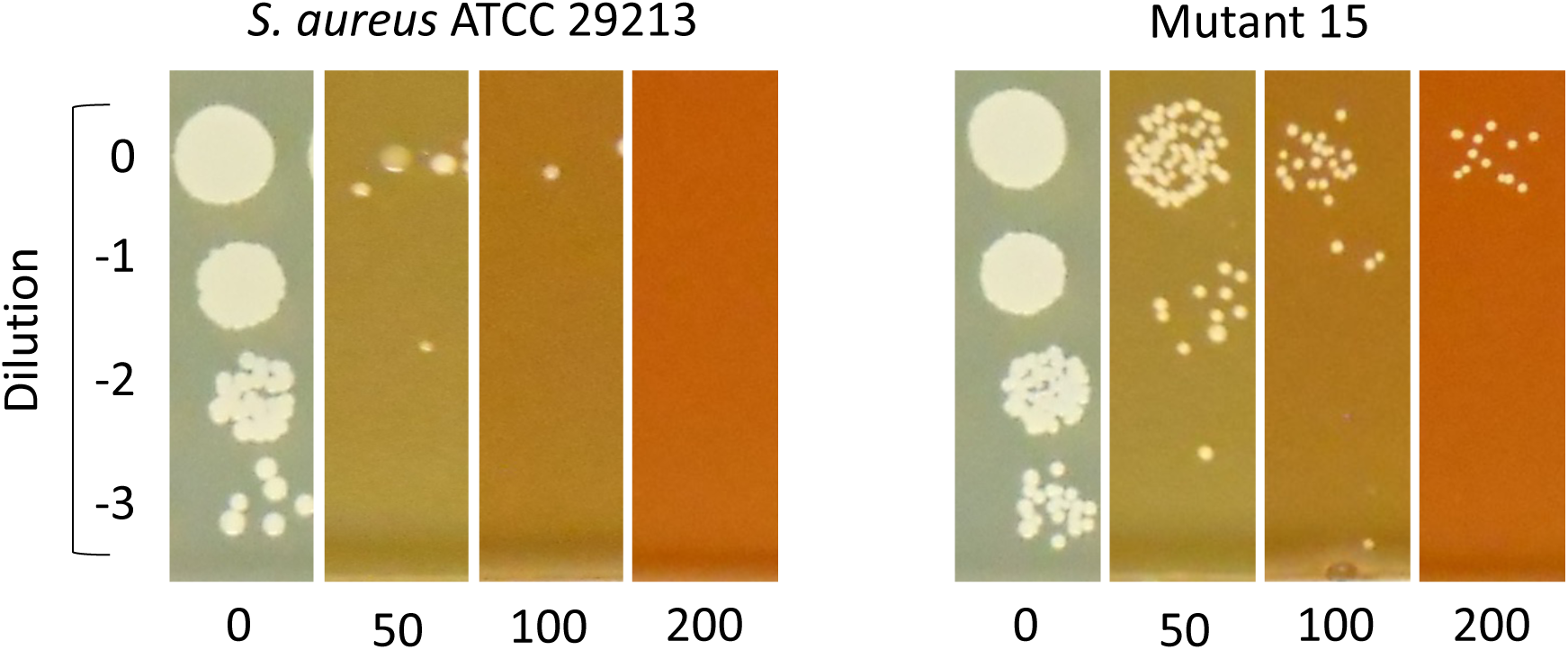
Drop dilution Compound 18 susceptibility testing of *S. aureus* mutants. Compound 18 concentrations are indicated in µM (horizontally).

We sequenced the genome of 17 mutants—15 from liquid cultures and 2 from agar plates—that were able to grow in the presence of 50-200 µM of Compound 18 better than the parental strain. The mutations identified, which were selected in two independent experiments, suggest a potential role of RNase Y in the susceptibility of *S. aureus* to Compound 18 (**Table S5)**. Among them, we validate genetically two nonsynonymous mutations (P280L and G240D) in the *rny* locus. Both engineered mutants displayed an increased MIC to Compound 18 (**Table 2**), confirming its role in *S. aureus* susceptibility.

**Table 2.** Antimicrobial activity of Compound 18 against genetically engineered *S. aureus rny* mutants.

| Strain | MIC (μM) |
| --- | --- |
| *S. aureus* ATCC 29213 | 6.25 |
| *S. aureus* ATCC 29213 Rny P280L | 50 |
| *S. aureus* ATCC 29213 Rny G240D | 25 |

We subsequently validated these results by time-kill kinetics. Surprisingly, we found that the kill curves of the engineered mutants were similar to that of the wild-type parental strain, with a rapid phase of initial killing followed by a rebound. However, the fraction of the population that survived this initial killing was approximately 10-fold higher in the mutant strains compared to the parental *S. aureus* strain, leading to a quicker recovery after the initial killing (**Fig. 4**).

**Fig. 4.**
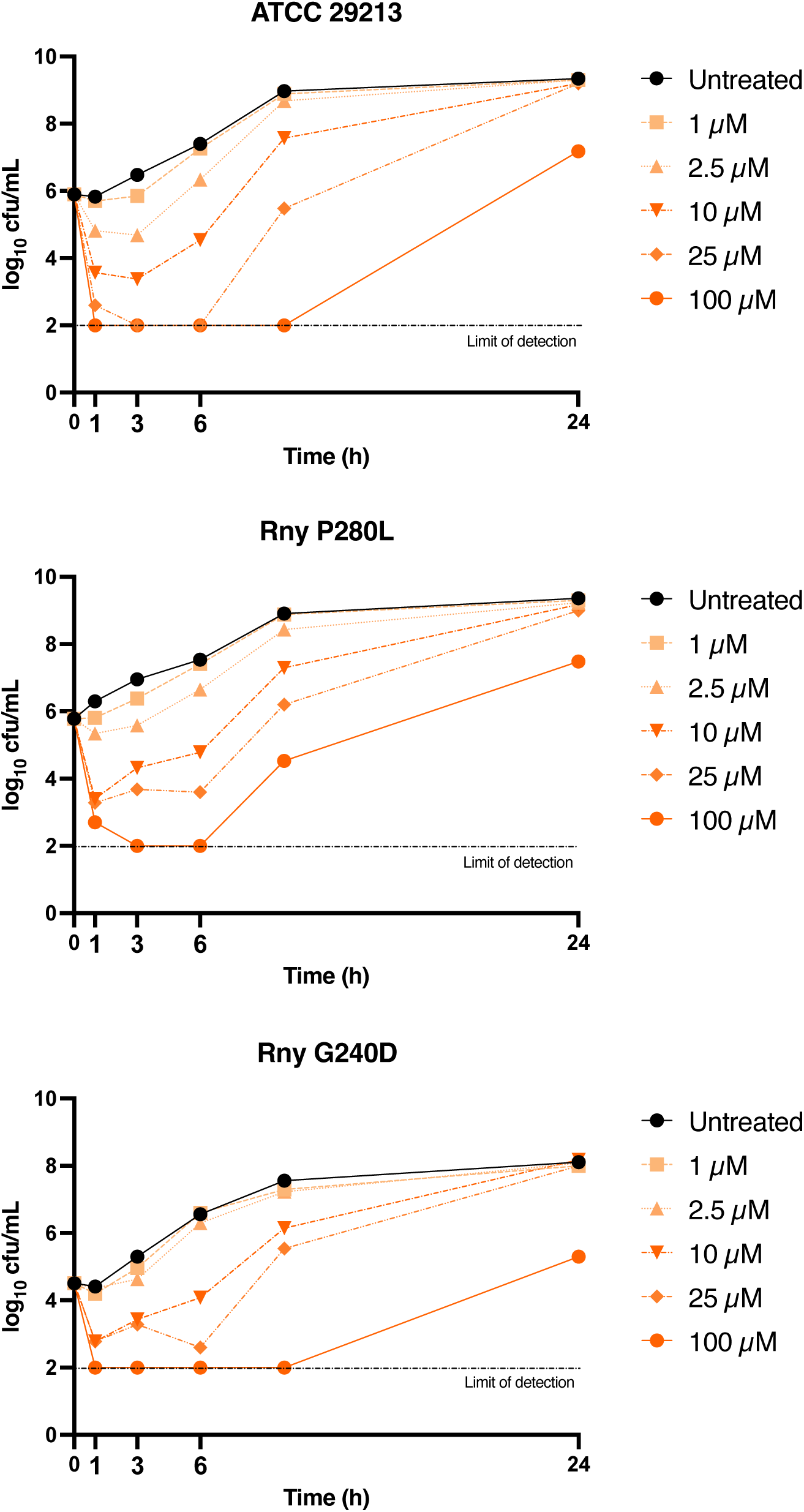
Time-kill kinetics of Compound 18 against *S. aureus rny* engineered point mutants. *S. aureus* ATCC 29213 (methicillin-susceptible, MIC = 25 µM) is included as the genetic background control.

### 4H-pyran-4-ylidene derivatives are not mutagenic

We tested the mutagenicity of the five compounds with broad-spectrum activity against Gram-positive bacteria (i.e., 02, 11, 18, 19, and 27) using the tryptophan reverse mutation assay. None of the compounds tested was mutagenic at 5 µM (0.2x MIC for *S. aureus* ATCC 29213) (**Fig. 5**). Since this assay is done by using *E. coli* and we previously observed a complete lack of activity of 4*H*-pyran-4-ylidene derivatives in Gram-negative bacteria (**Table S2**), we hypothesized that the absence of mutagenic activity of the compounds might be due to their limited penetration through the outer membrane, or to the activity of bacterial efflux pumps. Thus, we tested the activity of compounds 02, 18, and 19 against *E. coli* BW25113 mutant strains deficient in the Smr, EmrE, MdrA, and AcrB efflux pumps and combinations of them (Δ*emrE* Δ*mdrA*; Δ*emrE* Δ*acrB*; and Δ*emrE* Δ*mdrA* Δ*acrB*). We observed that the deletion of these efflux pumps did not increase the susceptibility of *E. coli* to 4*H*-pyran-4-ylidene derivatives at detectable levels, all mutant strains have the same MIC as the wild-type strain (MIC > 50 µM), which suggests that efflux is neither the cause of the intrinsic resistance to them in Gram-negatives nor the reason why the compounds are not mutagenic.

**Fig. 5.**
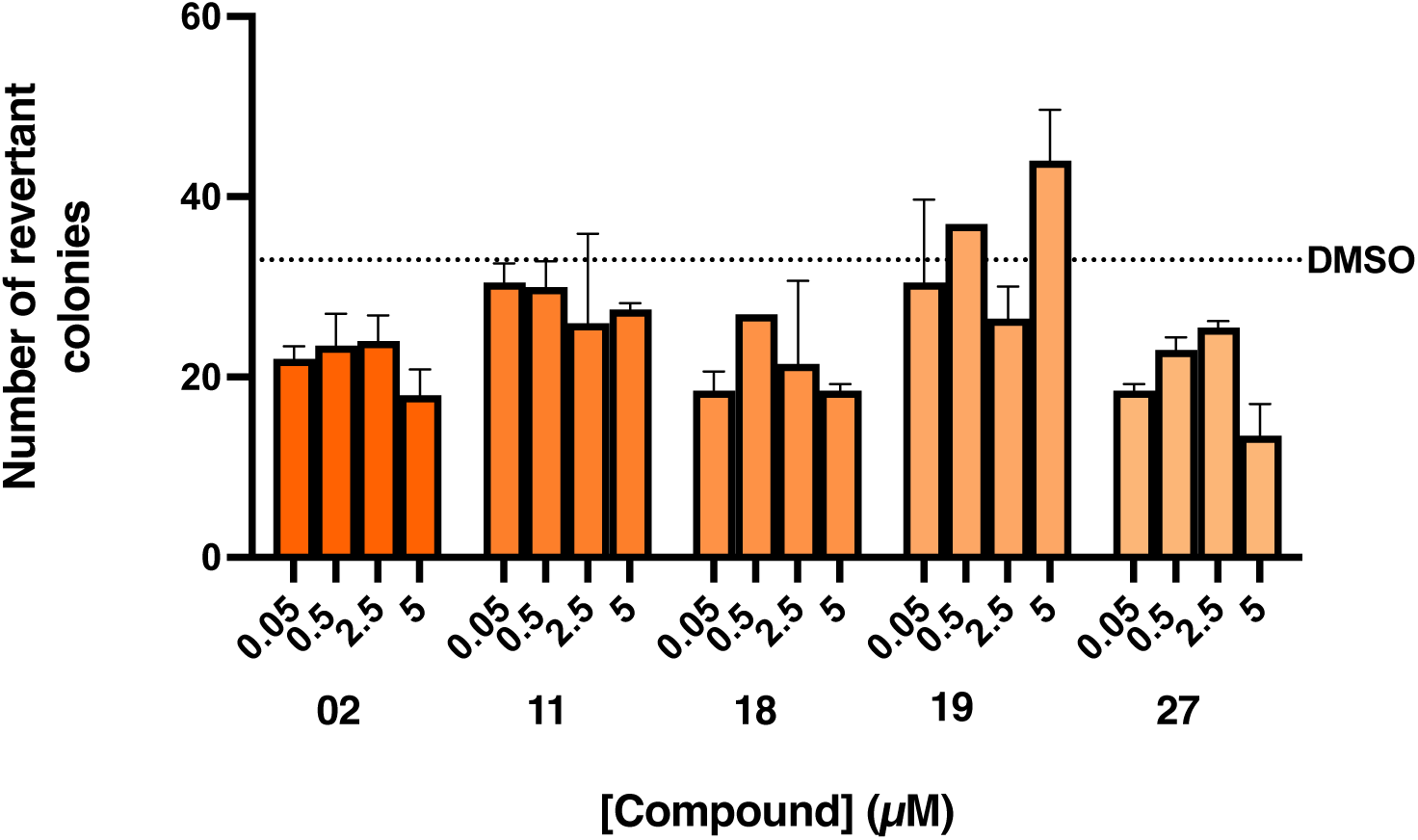
Tryptophan auxotrophy reversion test. Number of colonies isolated in the presence of each compound at the concentrations indicated horizontally. Compounds were considered mutagenic if the number of revertant colonies was twice the number in the DMSO control.

### 4H-pyran-4-ylidene derivatives are not cytotoxic to HepG2 cells

We determined the cytotoxicity of compounds 02, 11, 18, 19, and 27 against the HepG2 cell line using the neutral red assay. All five compounds were not cytotoxic at the concentrations at which antimicrobial activity was observed, with selectivity indexes ranging between 3.4 and 70 (**Table 3**).

**Table 3.** Cytotoxicity of 4H-pyran-4-ylidene derivatives against HepG2 cells and selectivity indices (SI, calculated as the ratio CC_50_/MIC_90_). CC, cytotoxic concentration.

| **Compound** | **CC (μM)** |  |  |  |  |
| --- | --- | --- | --- | --- | --- |
|  | **02** | **11** | **18** | **19** | **27** |
| $CC_{50}$ | 167 | 370 | 57 | 58 | 26 |
| $CC_{90}$ | 250 | >250 | 100 | 100 | 50 |
| **Strain** | **Selectivity index (SI, $CC_{50}/MIC_{90}$)** |  |  |  |  |
| *S. aureus* | 6.68 | 7.4 | 4.56 | <1 | <1 |
| *S. epidermidis* | 13.36 | 29.6 | 18.24 | 18.56 | <1 |
| *S. agalactiae* | 13.36 | 29.6 | 9.12 | 18.56 | 16.64 |
| *E. faecalis* | 6.68 | 7.4 | 36.48 | 74.26 | 8.32 |
| *C. glutamicum* | 26.72 | 14.8 | 9.12 | 18.56 | 4.16 |
| *C. diphtheriae* | 26.72 | 29.6 | 36.48 | 74.26 | 16.64 |

### Structure-activity relationships

After comparing data gathered from the dose-response assays with the structure of the compounds (**Table S2**), we observed some features related to their antimicrobial activity. To confirm these structure-activity relationships, we synthesized eight more derivatives and tested them in dose-response assays (compounds 40-47, **Table 4**). From these data, we inferred some features essential. First, *tert*-butyl substituents in the 4*H*-pyran-4-ylidene ring appeared to be necessary for antimicrobial activity, as their replacement by phenyl groups abolishes it. Second, carboxyl groups and the presence of a five-member heterocycle were also essential for activity of 4*H*-pyran-4-ylidene derivatives against Gram-positive bacteria; when removed, activity was lost. Bulky substituents, on the other hand, were not critical for antimicrobial activity (**Fig.6**).

**Fig. 6.**
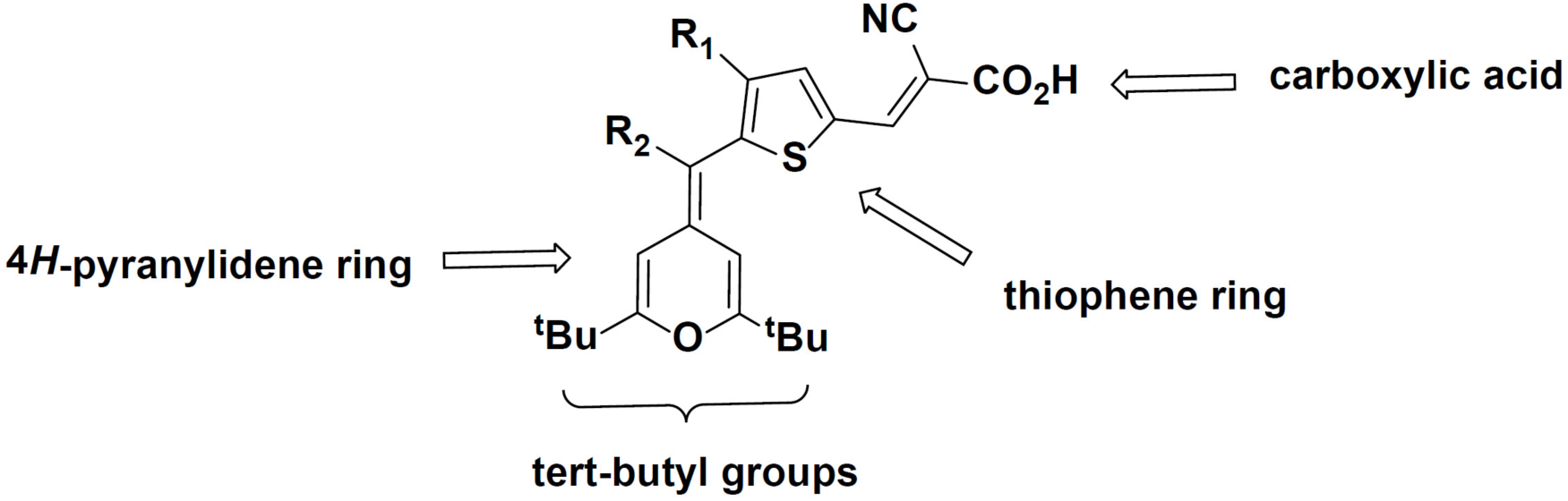
Pharmacophore of 4*H*-pyran-4-ylidenes. The presence of *tert-*butyl substituents, the pyranylidene and thiophene rings, and a carboxyl group are required for antimicrobial activity against Gram-positive bacteria.

**Table 4.** Antimicrobial activity of compounds 40-47. MIC, Minimal Inhibitory Concentration. MBC, Minimal Bactericidal Concentration. Values are given in µM. MIC/MBC values below 50 µM are indicated in yellow.

|  | *C. diphtheriae* |  | *C. glutamicum* |  | *E. faecalis* |  | *S. aureus*<br>(MSSA) |  | *S. epidermidis* |  | *S. agalactiae* |  |
| --- | --- | --- | --- | --- | --- | --- | --- | --- | --- | --- | --- | --- |
| *Compound* | MIC | MBC | MIC | MBC | MIC | MBC | MIC | MBC | MIC | MBC | MIC | MBC |
| 40 | >50 | >50 | >50 | >50 | >50 | >50 | >50 | >50 | >50 | >50 | >50 | >50 |
| 41 | >50 | >50 | >50 | >50 | >50 | >50 | >50 | >50 | >50 | >50 | >50 | >50 |
| 42 | 12.5 | 25 | >50 | >50 | >50 | >50 | >50 | >50 | >50 | >50 | >50 | >50 |
| 43 | 12.5 | 12.5 | 12.5 | 12.5 | 25 | 50 | 50 | >50 | 12.5 | 25 | 12.5 | 12.5 |
| 44 | 6.25 | 6.25 | 12.5 | 12.5 | 50 | 50 | 50 | 50 | 25-50 | 25-50 | 25 | 25 |
| 45 | 12.5 | 12.5 | 25 | 25 | 25 | 25 | 25 | 50 | 12.5 | 12.5 | 12.5 | 12.5 |
| 46 | 50 | 50 | >50 | >50 | >50 | >50 | >50 | >50 | >50 | >50 | 25 | 25 |
| 47 | 12.5 | 12.5 | 12.5 | 12.5 | 25 | 50 | 50 | 50 | 12.5 | 25 | 25 | 25 |

## Discussion

Only six new classes of antibacterial compounds have been introduced in the last five decades^16^. Several strategies have been proposed to broaden the pipeline of new antimicrobial candidates, including the use of assay conditions that mimic the *in vivo* environment in HTS campaigns, or the design of new chemical libraries based on the properties of compounds that are known to be able to penetrate bacteria^11,17^. Another approach to enrich this pipeline is searching for new antimicrobial natural products: the development of *in situ* culture techniques^18^, paired with the development of sequencing and bioinformatic tools, has allowed the identification of new antimicrobial candidates, such as teixobactin or darobactin^19,20^.

In this study, we took a different approach to seek for new antimicrobial scaffolds. For this, we screened an *in house* Photoactive Molecular Materials collection of compounds, originally designed to be part of solar cells, and identified a novel family with activity against Gram-positive bacteria, the 4*H*-pyran-4-ylidenes. This family showed no similarities with any compound available at the ZINC database when searching with the SwissSimilarity tool^21^. We then determined the activity of a panel of thirty-nine 4*H*-pyran-4-ylidene compounds, which were closely related to the initial hit that was identified in a single shot screening, and found that five of them were active against Gram-positive bacteria with MICs and MBCs ranging from 1.56 to 25 µM (**Table 1**).

We found that mutations in the *rny* locus, which encodes the RNase Y enzyme, decreased the susceptibility of *S. aureus* to Compound 18, one of the five 4*H*-pyran-4-ylidenes active against most of the Gram-positives tested. The RNase Y protein belongs to the RNA degradasome complex, which degrades RNA molecules and therefore determines their half-life. In addition to its ribonuclease activity, RNase Y also contains a transmembrane helix that anchors other ribonucleases belonging to the *S. aureus* degradasome^22^. The deletion of RNase Y in *S. aureus* stabilizes several tens of mRNAs, some of them encoding other components of the degradasome^23^. Given the killing kinetics observed for the mutants compared to the wild-type strain (**Fig. 4**), we hypothesize that mutations in *rny* are reducing the activity of the enzyme and consequently, changing RNA stability, which could increase the fraction of the population that is able to tolerate the initial killing induced by Compound 18. A similar phenomenon has been observed in *Mycobacterium tuberculosis*, in which the loss of RNase J—homologous to the RNase J1 and RNase J2 enzymes of the degradasome^24^—has been associated with tolerance to multiple drugs subsequent to the modification in the half-lives of multiple mRNAs^25^. The RNase Y enzyme is exclusive of Gram-positive bacteria, and its essentiality varies across species. In *Bacillus subtilis*, the most extensively studied Gram-positive species, the deletion of RNase Y results in slower growth and altered cell morphology. In *S. aureus*, on the other hand, the deletion of *rny* has no effect on bacterial growth, suggesting that RNase Y is not the direct target of 4*H*-pyran-4-ylidenes^24^.

There are two main possible explanations for the lack of activity of 4*H*-pyran-4-ylidenes against Gram-negative bacteria and mycobacteria. First, it is possible that their target is uniquely present in Gram-positive bacteria. Second, the more complex and less permeable cell walls present in mycobacteria and Gram-negative bacteria might hinder the access of these compounds to its intracellular target. Their lack of activity against *E. coli* mutants deficient in active transport proteins, suggests that efflux is not the reason for the intrinsic resistance, at least in Gram-negative species.

Preliminary toxicity studies in mammalian cell lines provided selectivity indices between 4.12 and 74. There is not a clear cutoff value to define a compound as safe based on its therapeutic index, but values higher than 10 are preferred in drug development programs^26^.

SAR studies showed that three substituents are needed for the activity of the 4*H*-pyran-4-ylidene family: the tert-butyl and carboxyl groups and a 5-member heterocycle. A better understanding of the mechanism of action will allow the refinement of the pharmacophore to achieve MIC values comparable to compounds currently used to treat *S. aureus* infections. We determined MICs in the low micromolar range for vancomycin (0.2 µM); cloxacillin (0.7 µM), and linezolid (1 µM).

*S. aureus* causes a wide array of diseases, from skin infections to fatal bacteremia. MRSA is currently the most abundant resistant pathogen in multiple regions of the world and is listed as a high priority for the development of novel antimicrobials. The main current therapeutic options for MRSA are vancomycin, daptomycin and linezolid, although the emergence of vancomycin-intermediate and resistant strains threaten its future availability as an option^15,27^. In the current scenario, exploring the chemical diversity intended for industrial indications could open a new path for the discovery of novel antimicrobials.

## Materials and methods

### Bacterial strains and culture conditions

Bacterial strains used in this study are listed in **Table S2**. Gram-positive and Gram-negative bacteria were propagated in Müller-Hinton broth (Panreac AppliChem) with 22 mg/L Ca^2+^ and 12 mg/L Mg^2+^ (Müller-Hinton II). *Corynebacterium diphtheriae* was grown in Brain Heart Infusion (BHI) broth (Difco). Mycobacterial strains were propagated in Middlebrook 7H9 broth (Difco) supplemented with 10% (v/v) Middlebrook ADC (Difco) and 0.05% Tween 80 (Scharlab).

### Compound management

Compounds were dissolved in DMSO at a final concentration of 4 mM and stored at –20°C. Secondary stocks were prepared in 96-well V-bottomed plates at 40-fold the final concentration in the assay plate.

### Screening of the Photoactive Molecular Materials (PMM) compound library

Antimicrobial activity was evaluated following a two-step process. First, a single-shot assay was performed with compounds being tested at a final concentration of 50 µM against strains described in **Table S2.** Compounds active at 50 µM were subsequently tested in dose-response assays in two-fold serial dilutions of the compounds, starting from 50 µM.

To perform each assay, mother plates were prepared with compounds at 40-fold the final concentration. Then, 5 µL were transferred onto 96-well flat-bottomed plates containing 195 µL of the corresponding bacterial suspension at a final density of 10^5^ CFU/mL. Plates were incubated for 24-72 hours (**Table S2**) before the addition of 30 µL of a solution containing Tween 80 (10%, v/v) and MTT [3-(4,5-dimethylthiazol-2-yl)-2,5-diphenyltetrazolium bromide] (2.5 mg/mL) (Sigma) as a reporter of bacterial growth. Optical density measures at 580 nm after 1-hour incubation were used to define the Minimal Inhibitory Concentration (MIC), which was the lowest compound concentration that inhibited 90% MTT conversion to formazan.

In order to determine the Minimal Bactericidal Concentration (MBC), 5 µL of each well were transferred to 96-well plates containing LB-agar (10 g/L tryptone, 5 g/L yeast extract, 5 g/L NaCl, and 17 g/L agar), before the addition of the MTT solution, and incubated for further 24 hours. The readout was done after the addition of 30 µL of a 0.1 mg/mL resazurin solution (Sigma) to each well and color changes from blue to pink were visually evaluated.

### Time-kill kinetics

A starting inoculum of 10^5^ CFU/mL of *S. aureus* was treated with 50, 25, 10, and 5 µM of Compound 18 (corresponding to 4, 2, 0.8, and 0.4-fold the MIC against *S. aureus* ATCC 29213). Then, 10-fold serial dilutions in PBS were seeded onto LB agar plates at 0, 1, 3, 6, and 24 hours, and CFUs enumerated after 24-hour incubation at 37°C.

### Mutant isolation assays

*S. aureus* ATCC 29213 cultures treated with 25 µM compound 18 for 24 hours were propagated in fresh Müller-Hinton II for an additional 24 hours. Then, cultures were treated with 100 µM of Compound 18 and incubated overnight before seeding 10-fold serial dilutions onto drug-free LB agar plates. Twenty-three colonies were selected for subsequent phenotypic validation.

Mutant isolation was also attempted by seeding 10^7^, 10^8^, and 10^9^ CFUs of *S. aureus* ATCC 29213 onto Müller-Hinton II agar plates with 100 µM of Compound 18. Plates were incubated at 37°C for 48 hours to allow selection of late-growing mutants. Eleven colonies were selected for phenotypic validation.

In both cases, the obtained colonies were transferred to Müller-Hinton II broth and grown cultures were used to determine the MIC of Compound 18 against them. Colonies with a 4-fold or greater increase in their MICs compared to the reference strain were selected for secondary validation, which consisted of seeding 5 µL of 10-fold serial dilutions on Müller-Hinton agar plates containing 50, 100, or 200 µM of Compound 18.

### Genomic DNA extraction

Mutants of *S. aureus* with decreased susceptibility to Compound 18 were grown overnight in tubes containing 5 mL of Müller-Hinton II broth. Bacteria were then collected by centrifugation and resuspended in 400 µL of TE buffer (10 mM Tris-HCl, 1 mM EDTA). Lysostaphin (Sigma) was then added at a final concentration of 0.1 mg/mL and the mixture was incubated for one hour at 37°C. Then 0.05 mg of proteinase K dissolved in 75 µL of a 10% sodium dodecyl sulfate solution were added and samples were incubated for 10 minutes at 65°C. Genomic DNA extraction was done by adding 750 µL of chloroform-isoamyl alcohol (24:1, v/v). Samples were mixed thoroughly before centrifugation (5 minutes, 13,500 RCF). The supernatant was transferred to tubes containing 420 µL of ice-cold isopropanol. DNA precipitation was carried out at –20°C for two hours before centrifuging samples (5 minutes, 13,500 RCF) to collect the nucleic acids. Pellets were finally resuspended in 50 µL of nuclease-free water (Qiagen). DNA was quantified by absorbance readings at 260 nm using an ND-1000 spectrophotometer (NanoDrop).

### Whole Genome Sequencing

Genomic DNA of seventeen mutants and the parental *S. aureus* ATCC 29213 was sequenced at the FISABIO Sequencing and Bioinformatics Service (Valencia, Spain) using Illumina technology or at the CIBA genomics facility (Zaragoza, Spain) using IonTorrent technology. Reads were filtered with fastp^42^ to remove low-quality bases at the 3’ end. The filtered reads were subsequently mapped to the *S. aureus* ATCC 29213 chromosome (available at https://genomes.atcc.org/genomes/21eea9803c88405a) with BWA^43^ and potential duplicates were removed with Picard tools (http://broadinstitute.github.io/picard). Single nucleotide polymorphisms (SNPs) were identified using VarScan^44^ if at least 20 reads supported the genomic position, the SNP was found at a frequency of 0.9 or higher and was not found near an indel region (10 bp). Indels were identified using Genome Analysis ToolKit (GATK)^45^. SNPs and indels were annotated using SnpEFF^46^, and those common to the parental strain were later removed.

### Genetic validation of mutations associated with Compound 18 resistance

Mutations identified in Compound 18-resistant *S. aureus* mutants were validated using the pMAD vector^47^ to perform allelic replacement in the parental strain. Mutant alleles were amplified by PCR using genomic DNA as the template and cloned between the *Xma*I and *Sfo*I restriction sites of pMAD. The resulting plasmids were then electroporated into the methylase-deficient *Escherichia coli* GM2929 to obtain plasmid DNA suitable for electroporation into *S. aureus*. One microgram of each plasmid was electroporated into electrocompetent *S. aureus* and transformants were selected on Tryptic Soy Agar (TSA, Difco) plates containing 2.5 µg/mL erythromycin and 80 µg/mL X-gal (5-Bromo-4-Chloro-3-Indolyl β-D-Galactopyranoside) after 72 hours of incubation at 28 °C. Then, one cyan colony from each transformation was transferred to Tryptic Soy Broth (TSB, Difco) with 2.5 µg/mL erythromycin and incubated at 44 °C overnight. This culture was inoculated onto TSA plates containing erythromycin and X-gal at 44°C to select colonies in which the recombinant pMAD derivative was integrated in the chromosome of *S. aureus*. Such cyan colonies were replicated onto TSA plates with erythromycin and X-gal and TSB without erythromycin. Liquid cultures grown overnight at 28°C were 10-fold serially diluted and seeded onto erythromycin-free TSA plates with X-gal. Plates were incubated at 37°C for 48 hours and white colonies (indicative of the loss of integrated pMAD derivative) were screened by colony PCR followed by Sanger sequencing to confirm the presence of the mutation.

### Cytotoxicity assays

Cytotoxicity of compounds 02, 11, 18, 19, and 27 (from a 10 mM DMSO stock solution) was tested in HepG2 cells (ECACC 85011430), which were exposed to a concentration of 250 µM and to a series of two-fold dilutions from 100 to 1.56 µM. Toxicity was determined using the Neutral Red Uptake (NRU) assay as described in ISO 10993-5:2009(E). Cells were seeded in 96-well flat-bottomed plates at a density of 2.5·10^4^ cells/well and incubated for 24 hours in Dulbecco Modified Eagle Medium (DMEM). Then, the culture medium was replaced with 100 µL of fresh DMEM with the compounds to be tested. The assay was performed in technical triplicates, including 2.5% DMSO (final concentration of DMSO in the 250 µM wells), 10% DMSO (death control), and 1 µg/mL rifampicin (positive control).

After incubation with the compounds for 24 hours, the culture medium was replaced by neutral red-containing medium. Cells were subsequently incubated for 3 hours before the desorb solution (1% glacial acetic acid, 50% ethanol, v/v) was added and optical density was measured at 540 nm. Compounds were also tested under the same conditions in plates without cells in order to ensure that their strong colors did not interfere with the readout method.

### Mutagenesis tests

*Escherichia coli* WP2 (*uvrAB trpE(AT), pKM101*) was inoculated on minimal agar medium with compounds 02, 11, 18, 19 and 27 at concentrations of 5, 2.5, 0.5 and 0.05 µM. Methylmethanesulfonate, a well-known mutagenic compound was included as positive control for mutagenesis. Plates were incubated for 72 h at 37°C and colonies were subsequently counted. All determinations were performed in duplicates.

### Compound synthesis and characterization

#### Scheme synthesis of compound (03): 2-cyano-3-(7-((2,6-diphenyl-4*H*-pyran-4-ylidene)methyl)-2,3-dihydrothieno[3,4-b][1,4]dioxin-5-yl)acrylic acid

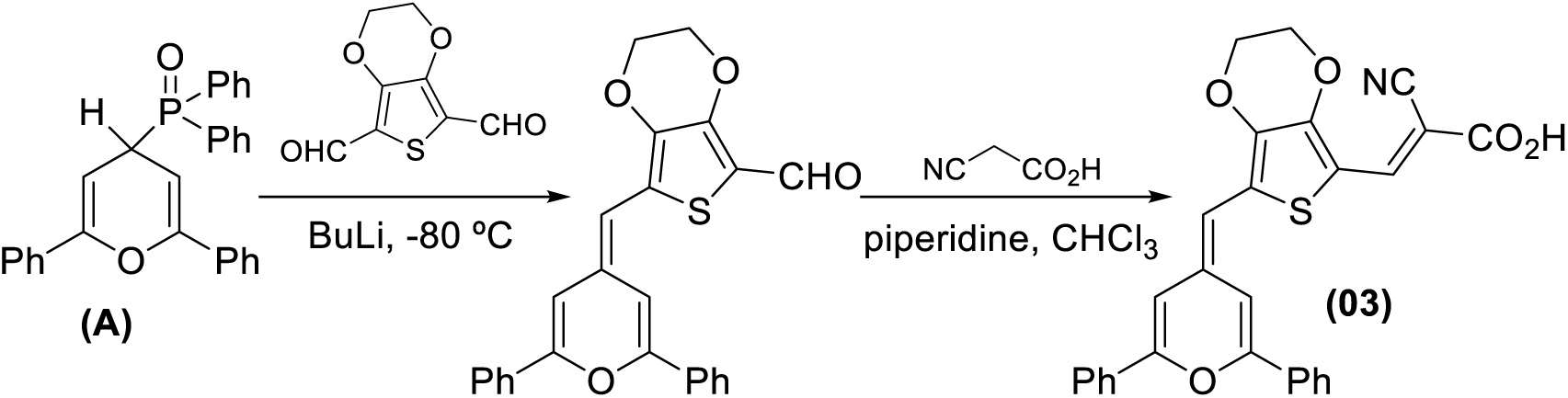

Step 1: A solution of (2,6-diphenyl-4*H*-pyran-4-yl)diphenylphosphine oxide (**A**)^28^ (1.11g, 2.55 mmol) in anhydrous THF (30 mL) was cooled to −80°C under an Ar atmosphere. Subsequently, *n*-BuLi (1.6 M in hexanes, 1.73 mL, 2.77 mmol) was added, and the mixture was stirred for 15 min at −78 °C. After this, a solution of 3,4-ethylenedioxythiophene-2,5-dicarboxaldehyde (commercially available) (502 mg, 2.53 mmol) in anhydrous THF (20 mL) was added, allowing the temperature to gradually return to room temperature over 18 hours. The reaction was then quenched with 5 mL of saturated NH_4_Cl and stirred for an additional 30 minutes. The product was extracted with AcOEt (3 × 20 mL), the organic phase was dried over anhydrous MgSO_4_, and the solvent was removed under reduced pressure. Purification by column chromatography using hexane/CH_2_Cl_2_ (98:2) as the eluent yielded the desired aldehyde as an orange solid (616 mg, yield: 59%).

Step 2: To a solution of the aldehyde prepared in step 1 (151 mg, 0,33 mmol) and 2-cyanoacetic acid (47 mg, 0.56 mmol) in anhydrous chloroform (7 mL) piperidine (240 μl) was added. The mixture was refluxed for 24 hours under an Ar atmosphere, then cooled to room temperature and acidified to a pH ≈ 2 of with a solution of HCl (1N). The organic phase was washed with water, and the solvent was evaporated. The residue was washed with a hexane/CH_2_Cl_2_ (90:10) mixture and dried to yield a dark violet solid (143 mg, 91%).

**M.p.:** 255-262 ^°^C. IR (KBr) cm^-1^: 3675-3135 (COOH), 2204 (C≡N), 1682 (C=C).

**^1^H NMR** (300 MHz, dmso-d_6_) 8 (ppm): 8,11 (s, 1H), 8,00−7,80 (m, 4H), 7,61−7,44 (m, 6H), 7,23 (brs, 1H), 7,15 (brs, 1H), 6,26 (s, 1H), 4,52−4,42 (m, 2H), 4,41−4,32 (m, 2H).

**HRMS (ESI^+^)** m/z: Calculated C_28_H_19_NO_5_S: 481.0978. Found: 481.0964

#### Scheme synthesis of Compound (06): 2-((5-((2,6-diphenyl-4*H*-pyran-4-ylidene)methyl)thiophen-2-yl) methylene)malononitrile

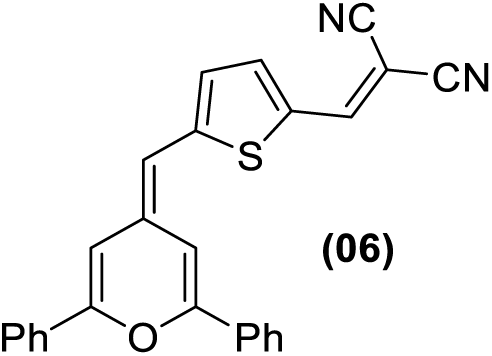

The starting aldehyde was synthesized in high yield according to a previously described procedure^13^. Piperidine (100 μl) was added to a solution containing the aldehyde precursor (73 mg, 0.20 mmol) and malononitrile (15.9 mg, 0.24 mmol) in 10 mL of ethanol. The reaction mixture was refluxed for 6 hours under an Ar atmosphere and then cooled in an ice bath. The resulting solid was filtered, washed with cold ethanol, and subsequently with a hexane/dichloromethane mixture (95:5). The residue was purified by flash chromatography (using hexane/CH_2_Cl_2_, 4:6 as the eluent) to afford an intense bluish-green solid (65 mg, 85%).

**M.p.:** 258-260 ^°^C. IR (Nujol) cm^-1^: 1608 (C=N).

**^1^H NMR** (400 MHz, CDCl_3_) d (ppm): 8.72 (d, *J* = 12.2 Hz, 1H), 7.28 (d, *J* = 1.8 Hz, 1H), 6.41 (d, *J* = 1.8 Hz, 1H), 5.42 (d, *J* = 12.2 Hz, 1H), 3.56 (s, 3H), 3,18 (s, 3H), 1.37 (s, 9H), 1.31 (s, 9H).

**^13^C NMR** (100 MHz, DMSO-d6) 8 (ppm): 174.4, 174.1, 159.3, 158.1, 108.4, 102.7, 98.5, 47.1, 38.7, 37.1, 36.7, 27.9, 27.7

**HRMS (ESI+)** m/z: Calculated C_26_H_17_N_2_OS: 405.1056. Found: 405.1044

#### Scheme synthesis of Compound (07): ethyl hydrogen-(1-cyano-2-(5-((2,6-diphenyl-4*H*-pyran-4-ylidene)methyl)thiophen-2-yl)vinyl)phosphonate

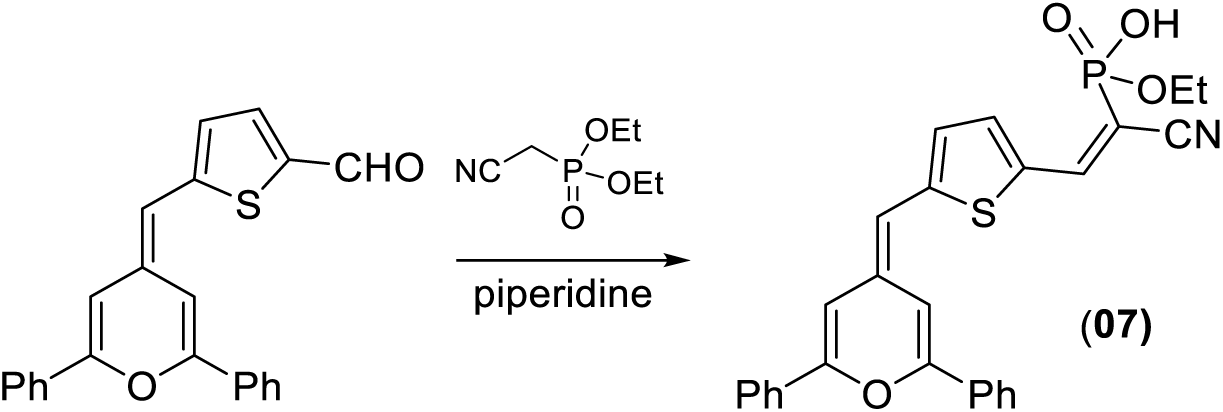

The starting aldehyde was synthesized in high yield according to a previously described procedure^13^. Piperidine (400 μl) was added to a solution of aldehyde precursor (206 mg, 0.578 mmol) and diethyl cyanomethylphosphonate (114 μl, 0.69 mmol) in 10 mL of anhydrous acetonitrile. The reaction mixture was heated to reflux under an Ar atmosphere for 24 hours. After solvent removal, the residue was purified by reversed-phase C18 chromatography (CH_3_CN/AcONH_4_ 20 mM 7:3 as eluent) to give a maroon solid (153 mg, 54%) as a Z/E mixture.

**M.p.:** 200-203 ^°^C.

**^1^H NMR** (400 MHz, DMSO-d6) 8 (ppm): 8.11-7.84 (m, 5H), 7.81 (d, *J* = 4.1 Hz, 1H), 7.63-7.44 (m, 6H), 7.28 (br s, 1H), 7.22 (d, *J* = 4.1 Hz, 1H), 7.02 (br s, 1H), 6.36 (s, 1H), 4.04-3.89 (m, 2H), 1.25 (t, *J* = 7.0 Hz, 3H).

**^13^C NMR** (100 MHz, DMSO-d6) 8 (ppm): 154.2, 151.6, 149.7, 147.5, 147.4, 138.5 (x2), 133.4, 133.2, 131.8, 131.5, 130.5, 129.8, 129.1, 128.9, 126.3, 124.9, 124.5, 117.9, 117.8, 108.6, 107.3, 102.3, 95.3, 93.4, 61.5, 61.4, 16.3, 16.2

**HRMS (ESI^+^)** m/z: Calculated C_27_H_23_NO_4_PS: 488.1079. Found: 488.1080

#### Scheme synthesis of Compound (10): (1-cyano-2-(2-((2,6-di-tert-butyl-4H-pyran-4-ylidene)methyl) thiazol-5-yl)vinyl)phosphonic acid

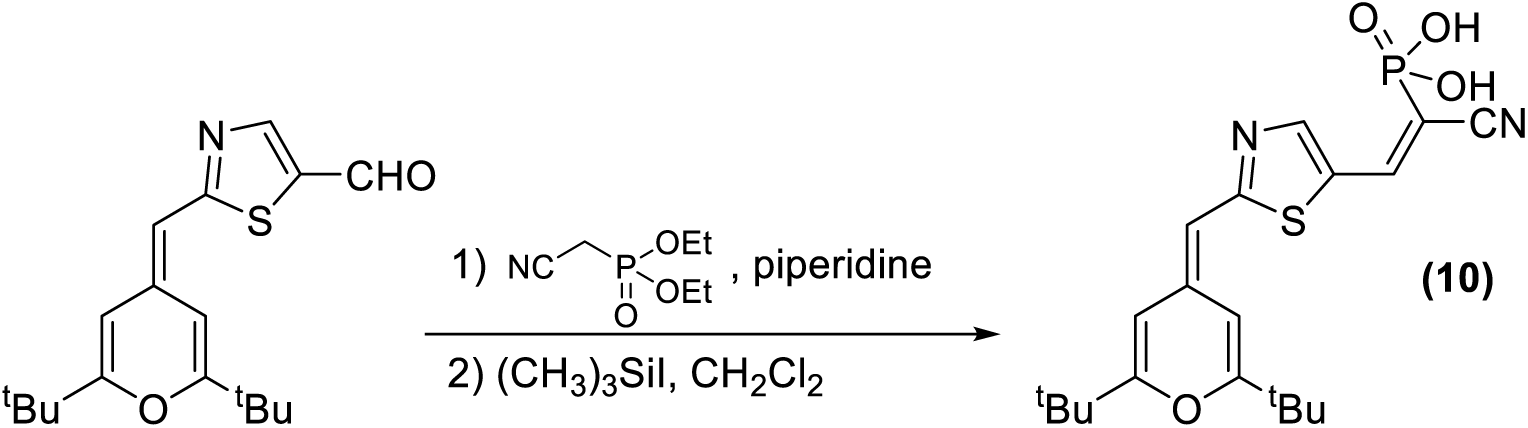

The starting aldehyde was synthesized in high yield according to a procedure previously described^29^. Piperidine (250 ml) was added to a solution of aldehyde precursor (131 mg, 0.356 mmol) and diethyl cyanomethylphosphonate (71 ml, 0.428 mmol) in 8 mL of anhydrous acetonitrile. The reaction mixture was heated to reflux for 24 hours under an Ar atmosphere. After removing the solvent, the residue was purified by flash chromatography (CH_2_Cl_2_/AcOEt, 10:1 as eluent) to give a red oil (142 mg, 85%).

Subsequently, trimethylsilyl iodide (206 μl, 1.397 mmol) was added to a solution of the intermediate obtained in the previous step (108 mg, 0.226 mmol) in anhydrous dichloromethane (15 mL). The resulting mixture was refluxed for 2 hours. Subsequently, 10 mL of methanol was added, and the mixture was stirred at room temperature for 1 hour. After solvent evaporation, the residue was washed with a hexane/CH_2_Cl_2_ mixture (9:1) until the washing solution became colorless. The resulting residue was dried under vacuum to yield a maroon solid (85 mg, 89%).

**M.p.:** 193-196 ^°^C.

**^1^H NMR** (400 MHz, DMSO-d6) 8 (ppm): 8.28 (s, 1H), 7.98 (s, 1H), 7.92 (s, 1H), 7.52 (br s, 1H), 6.14 (br s, 1H), 6.07 (s, 1H), 1.22 (s, 9H), 1.19 (s, 9H).

**HRMS (ESI^+^)** m/z: Calculated C_20_H_25_N_2_NaO_4_PS: 443.1170. Found: 443.1159

#### Scheme synthesis of Compound (13): 5-(2-(2,6-di-*tert*-butyl-4*H*-pyran-4-ylidene)ethylidene)-4-(4-methoxyphenyl)-2-oxo-2,5-dihydrofuran-3-carbonitrile

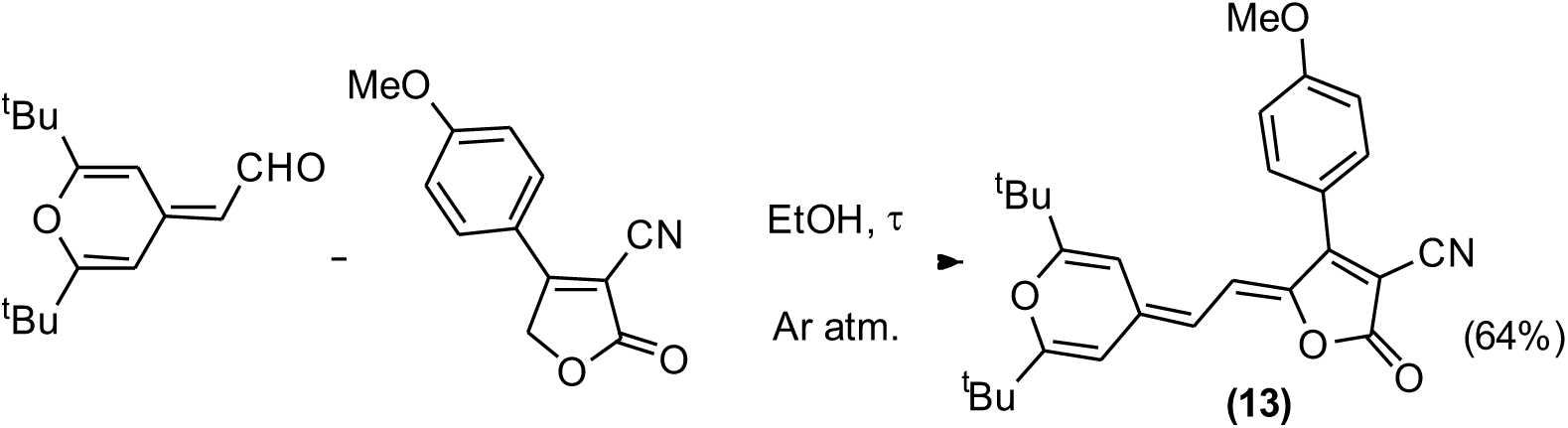

To a solution of 2,6-di-*tert*-butyl-4-formylmethylene-4*H*-pyran^30^ (141 mg, 0.60 mmol) in absolute ethanol (5 mL), 4-(4-methoxyphenyl)-2-oxo-2,5-dihydrofuran-3-carbonitrile^31^ (129 mg, 0.60 mmol) was added. The mixture was refluxed under argon with exclusion of light for 16.5 hours. After cooling, the resulting solid was isolated by filtration, washed with cold ethanol, and then with mixture of cold pentane/CH_2_Cl_2_ 9.5:0.5, affording a dark blue solid (166,3 mg; yield: 64%).

**M.p.:** 197−200 ^°^C. **IR** (nujol): ρ̅ (cm^-1^) 2212 (C≡N), 1739 (C=O), 1654 (C=C), 1609 and 1583 (C=C, Ar).

**^1^H NMR** (400 MHz, CDCl_3_) δ (ppm): 7.60−7.55 (m, 2H), 7.09−7.04 (m, 2H), 6.67 (d, *J* = 13.0 Hz, 1H), 6.18 (d, *J* = 1.8 Hz, 1H), 6.11 (d, *J* = 13.0 Hz, 1H), 6.04 (d, *J* = 1.8 Hz, 1H), 3.90 (s, 3H), 1.25 (s, 9H), 1.24 (s, 9H).

**^13^C NMR** (100 MHz, CDCl_3_): δ (ppm) 169.3, 168.9, 165.8, 162.1, 157.5, 144.6, 141.5, 130.6, 121.4, 120.4, 114.7, 114.0, 108.3, 107.3, 100.1, 90.3, 55.6, 36.3, 36.0, 27.8, 27.5.

#### Scheme synthesis of compound (14): 2-cyano-3-(7-(4-((2,6-diphenyl-4*H*-pyran-4-ylidene)methyl)phenyl) benzo[c][1,2,5]thiadiazol-4-yl)acrylic acid

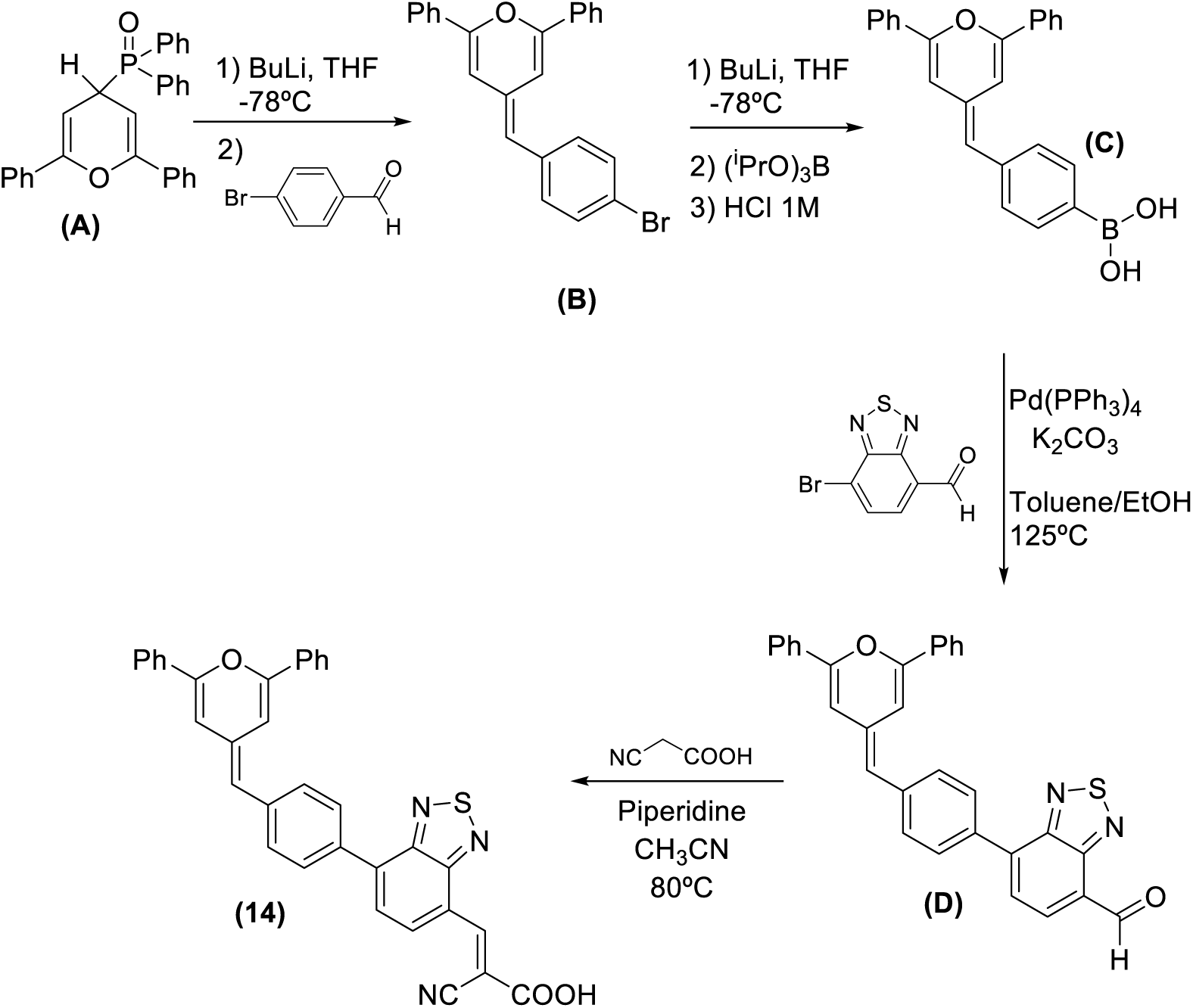

#### Compound (B): 4-(4-bromobenzylidene)-2,6-diphenyl-4*H*-pyran

A solution of (2,6-diphenyl-4*H*-pyran-4-yl)diphenylphosphine oxide **A**^28^ (547 mg, 1.27 mmol) in anhydrous THF (10 mL) was cooled to −78°C under an Ar atmosphere. Subsequently, n-BuLi (1.6 M in hexanes, 0.86 mL, 1.4 mmol) was added, and the solution was stirred for 20 min at −78 °C. Afterward, a solution of *p*-bromobenzaldehyde (commercially available) (234.8 mg, 1.27 mmol) in anhydrous THF (10 mL) was added, allowing the temperature to gradually return to room temperature over 18 hours. The reaction was quenched with 5 mL of a saturated NH_4_Cl solution and stirred for an additional 30 minutes. The product was extracted with CH_2_Cl_2_ (2 × 15 mL), the organic phase was dried over anhydrous MgSO_4,_, and the solvent was removed under reduced pressure. The product was purified by column chromatography with hexane/CH_2_Cl_2_ (9:1) as the eluent, yielding the desired product as an orange solid (367 mg, yield: 74%).

**M.p.:** (°C): 194−196. IR (KBr): ρ̅ (cm^-1^) 1657, 1600, 1579 and 1545 (C=C, Ar), 1075 (Csp^2^-Br).

**^1^H NMR** (300 MHz, CD_2_Cl_2_) δ (ppm): 7.82−7.76 (m, 4H), 7.50−7.41 (m, 8H), 7.32−7.26 (m, 2H), 6.95 (dd, *J*_1_ = 1.8 Hz, *J*_2_ = 0.6 Hz, 1H), 6.45 (d, *J* = 1.8 Hz, 1H), 5.88 (s, 1H).

**^13^C NMR** (100 MHz, CDCl_3_): δ (ppm): 153.6, 151.5, 138.1, 133.8, 133.6, 132.0, 130.5, 130.0, 129.9, 129.7, 129.2, 125.4, 125.0, 119.2, 113.3, 108.8, 102.1.

**HRMS (ESI^+^)**: m/z calculated for C_24_H_18_BrO ([M+H]^+^) 401.0536, found 401.0546.

#### Compound (C): 4-((2,6-diphenyl-4*H*-pyran-4-ylidene)methyl)phenylboronic acid

A solution of compound **B** (243 mg, 0.67 mmol) in anhydrous THF (8 mL) was cooled at −78°C under an Ar atmosphere. Subsequently, *n*-BuLi (1.6 M in hexanes, 0.54 mL, 0.73 mmol) was added, and the solution was stirred for 1h to −78 °C. Afterwards, triisopropyl borate (0.21 mL, 0.92 mmol) was added, and the temperature was allowed to gradually return to room temperature over 22 hours. Next, 6 mL of HCl (1N) were added and the mixture was stirred for 30 min. The product was extracted with CH_2_Cl_2_ (2 × 20 mL), the organic phase was dried over anhydrous MgSO_4,_ and the solvent was removed under reduced pressure. To the crude oil, 10 mL of ethyl acetate were added, resulting in a yellow solid, which was then filtered and washed with cold ethyl acetate, yielding a yellow solid (68.4 mg, yield: 28%).

**M.p.:** (°C): 217−220. IR (KBr): ρ̅ (cm^-1^) 3243 (BO-H), 1658 (C=C, Ar), 1591 and 1572 (C=C, Ar), 1376 (B-O).

**^1^H NMR** (400 MHz, THF-d_8_) δ (ppm): 7.84−7.77 (m, 4H), 7.75 (d, *J* = 8.1 Hz, 2H), 7.45−7.33 (m, 8H), 7.08 (d, *J* = 1.8 Hz, 1H), 7.04 (s, 2H), 6.59 (d, *J* = 1.8 Hz, 1H), 5.97 (s, 1H).

**^13^C NMR** (100 MHz, THF-d_8_) δ (ppm): 154.0, 152.0, 141.6, 135.6, 135.0, 134.8, 130.8, 130.7, 130.6, 130.3, 130.0, 129.9, 128.1, 126.2, 125.8, 116.4, 110.0, 103.5.

**HRMS (ESI^+^)**: m/z calculated for C_24_H_19_BO_3_ ([M]^+·^) 366.1426, found 366.1449.

#### Compound (D): 7-(4-((2,6-diphenyl-4*H*-pyran-4-ylidene)methyl)phenyl)benzo[*c*] [1,2,5]thiadiazole-4-carbaldehyde

A solution of boronic acid **C** (69 mg, 0.189 mmol), 7-bromobenzo[c][1,2,5]thiadiazole-4-carbaldehyde^32^. (33.5 mg, 0.164 mmol), K_2_CO_3_ (239 mg, 1.64 mmol) and tetrakis (triphenylphosphine)palladium(0) (19.1 mg, 0.016 mmol) in 25 mL of toluene and 5 mL of ethanol, previously deoxygenated, was heated at 125 °C for 45 min. After cooling, the solvent was evaporated and the product was purified by column chromatography using hexane/CH_2_Cl_2_ (1:9) as eluent, yielding a dark red-purple solid (55 mg, yield: 83%).

**M.p.:** (°C): 188−190. IR (KBr): ρ̅ (cm-1) 2921 and 2848 (C−H), 1680 (C=O), 1656 (C=N), 1583 and 1535 (C=C, Ar).

**^1^H NMR** (400 MHz, CD2Cl2) δ (ppm): 10.78 (s, 1H), 8.30 (d, J = 7.4 Hz, 1H), 8.09−8.06 (m, 2H), 7.94 (d, J = 7.4 Hz, 1H), 7.85−7.80 (m, 4H), 7.62−7.60 (m, 2H), 7.51−7.40 (m, 6H), 7.15 (d, J = 1.9 Hz, 1H), 6.53 (d, J = 1.9 Hz, 1H), 6.04 (s, 1H).

**HRMS (ESI^+^):** m/z calculated for C_31_H_20_N_2_NaO_2_S ([M+Na]^+^) 507.1138, found 507.1149.

#### Scheme synthesis of Compound 14: 2-cyano-3-(7-(4-((2,6-diphenyl-4*H*-pyran-4-ylidene)methyl)phenyl)benzo[c][1,2,5]thiadiazole-4-yl)acrylic acid (14)

A solution of aldehyde **D** (46 mg, 0.095 mmol), cyanoacetic acid (12 mg, 0.142 mmol), and piperidine (62 µL; 0.627 mmol) in dried acetonitrile (6 mL) was heated at 85 °C (TLC monitoring; eluent: hexane/CH_2_Cl_2_ (1:9)) for 24 hours. After cooling, the resulting solid was isolated by centrifugation at 40,000 rpm. The product was then purified by washing with a mixture of hexane/CH_2_Cl_2_ (9:1). The solid was recovered by centrifugation (40,000 rpm), yielding a dark red solid. (46 mg, yield: 89%).

**M.p.:** (°C): 219−221. IR (KBr): ρ̅ (cm^-1^) 3433 (O-H, broad), 2213 (C≡N), 1655 (C=O), 1638 (C=N), 1577 (C=C, Ar), 1529 (C=C, Ar), 1337 (C-O).

**^1^H NMR** (400 MHz, DMSO-d_6_) δ (ppm): 8.74 (s, 1H), 8.59 (d, *J* = 7.5 Hz, 1H), 8.13 (d, *J* = 8.3 Hz, 2H), 8.10 (d, *J* = 7.5 Hz, 1H), 7.92−7.88 (m, 2H), 7.64 (d, *J* = 8.3 Hz, 2H), 7.56−7.44 (m, 6H), 7,16 (br s, 1H), 6.88 (d, *J* = 1.2 Hz, 1H), 6.12 (s, 1H).

**^13^C NMR** (100 MHz, DMSO-d_6_) δ (ppm): 162.4, 154.7, 152.9, 152.7, 150.3, 140.6, 138.9, 134.8, 133.5, 132.8, 132.6, 130.1, 129.9, 129.8, 129.7, 129.3, 129.2, 128.3, 128.1, 127.6, 125.2, 125.1, 124.6, 119.6, 114.5, 109.2, 102.0. HRMS (ESI^-^): m/z calculated for C_33_H_20_N_3_OS ([M-COOH]^-^) 506.1333, found 506.1328.

#### Scheme synthesis of Compound 20 and Compound 21

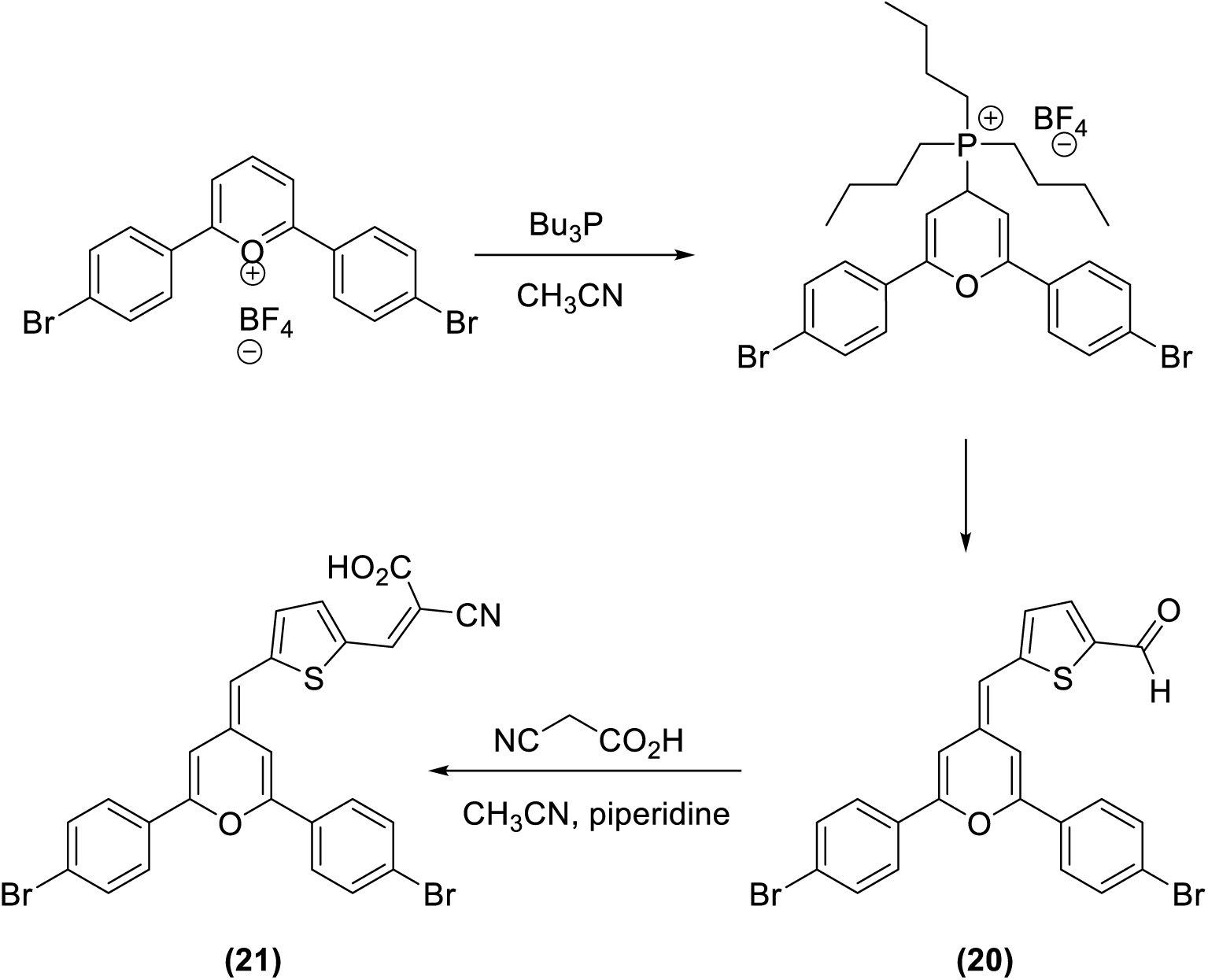

#### 4’-Bromo-2,6-diphenylpyrylium tetrafluoroborate

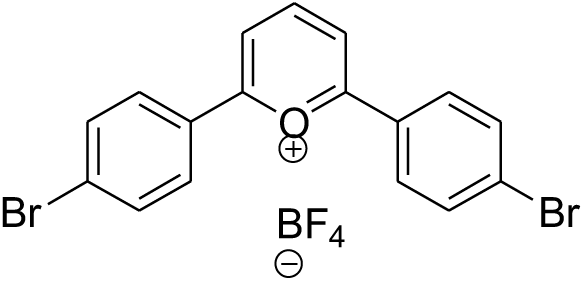

To a solution of ethyl orthoformate (5 mL, 30 mmol) and 2-bromo-1-phenylethan-1-one (1.998 g, 10 mmol), HBF_4_ (0.5 mL) and acetic anhydride (5 mL) were added at 0 °C. Then, the mixture was stirred for 90 minutres at room temperature and then to 50 °C for 30 minutes. The reaction was cooled to room temperature and the yellow solid filtered and washed with cood ethyl ether. Yield: 1.077 g, 45%

**M.p.:** 225 – 230 °C. **IR** (KBr) cm^-1^: 3098,51 (Csp^2^-H), 1504,33 (C=C).

**^1^H NMR** (300 MHz, acetone-d6) δ (ppm): 9.34 (dd, J_1_ = 8.0 Hz, J_2_ = 8.6 Hz, 1H), 8.98 (d, J = 8.3 Hz, 2H), 8.5 (dt, J_1_ = 9.0 Hz, J_2_ = 2.4 Hz, 4H), 8.0 (dt, J_1_ = 8.7 Hz, J_2_ = 2.7 Hz, 4H)

**13C NMR:** The product is unstable in deuterated solvents and poorly soluble. Its

^13^C-NMR spectrum could not be determined.

#### Tetrafluoroborate of tributyl(2,6-bis(4-bromophenyl)-4*H*-pyran-4-yl) phosphonium

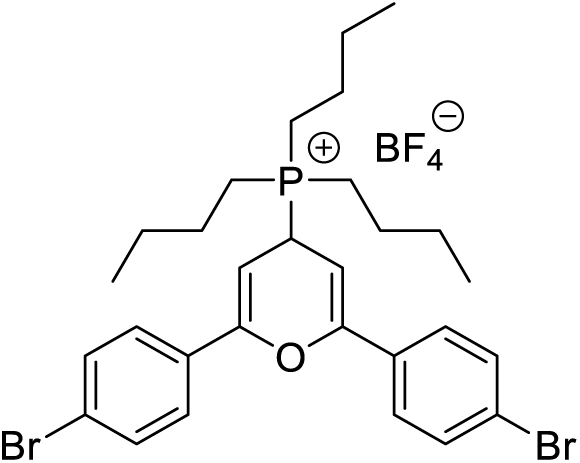

To a solution of pyrilium salt obtained in step 1 (0.573 g, 1.20 mmol) in anhydrous acetonitrile (3.5 mL) *n*-tributylphosphine (0.31 mL, 1.24 mmol) was added and the mixture was stirred for 2 h at room temperature. Then, 150 mL of cold ethyl ether were added and the solid was filtered and washed with cold ethyl ether. Yield: 749 mg, 92%, as an ochre-colored solid.

**M.p.:** 192 – 200 °C**. IR** (KBr) cm^-1^: 2958,54 (Csp^3^-H), 1488,65 (C=C).

**^1^H NMR** (400 MHz, Acetone-d6) δ (ppm): 7.81 (dt, J_1_ = 8.4 Hz, J_2_ = 1.8 Hz, 4H), 7.67 (dt, J_1_ = 8.8 Hz, J_2_ = 2.2 Hz, 4H), 5,97 (dd, J_1_ = 2.90 Hz, J_2_ = 4.9 Hz, 2H), 4.7 (dt, J_1_ = 14.4 Hz, J_2_ = 5.2 Hz, 1H) 2.61 (m, 6H), 1.80 (m, 6H), 1.51 (m, 6H), 0.92 (t, J = 7.3 Hz, 9H).

**^13^C NMR** (75 MHz, Acetone-d6) δ (ppm): 154.4, 154.3, 133.6, 128.8, 125.3, 92.6, 92.5, 32.0, 31.4, 25.7, 25.5, 25.2, 25.1, 18.4, 17.8, 14.6.

**HRMS (ESI^+^)** m/z: Calculated for C_29_H_38_Br_2_OP [M]: 591.1022. Found: 591.1052.

#### 5-((2,6-Bis(4-bromophenyl)-4H-pyran-4-ylidene)methyl)thiophene-2-carbaldehyde

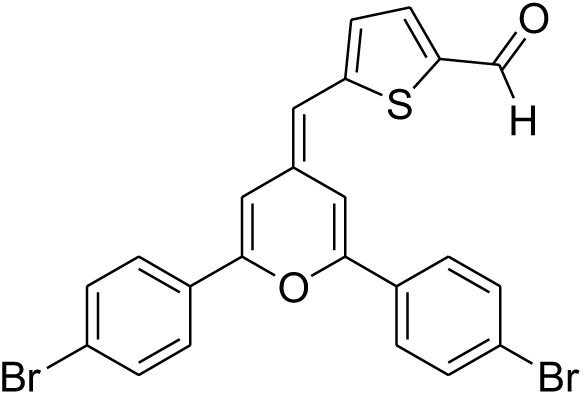

To a solution of the phosphonium salt (250 mg, 0.368 mmol) and potassium tert-butoxide (49.1 mg, 0.438 mmol) in anhydrous THF (15 mL) and the mixture was stirred for 15 minutes. Then thiophene-2,5-dicarbaldehyde (61 mg, 0.436 mmol) in THF (5 mL) was added and the mixture of reaction was stirred for 24 h. The reaction was quenched with a saturated solution of NH_4_Cl and extracted with ethyl acetate. The organic layer was dried over MgSO4 and purified by silica gel column chromatography (20% ethyl acetate in hexanes). Yield: 144 mg, 76% as a reddish solid.

**M.p.:** 209 – 214 ⁰C. **IR** (KBr) cm^-1^: 1652,42 (C=O), 1558,45 (C=C).

**^1^H NMR** (400 MHz, CH_2_Cl_2_) δ (ppm): 9.85 (s, 1H), 7.78 (dt, J_1_ = 8.8 Hz, J_2_ = 2.2 Hz, 2H), 7.67 (m, 7H), 7.25 (dd, J_1_ = 0.65 Hz, J_2_ = 1.97 Hz, 1H), 7.07 (dd, J_1_ = 0.58 Hz, J_2_ = 4.05 Hz, 1H), 6.58 (d, J = 1.62, 1H), 6.25 (s, 1H).

**^13^C NMR** (100 MHz, CH_2_Cl_2_): 182.5, 154.4, 152.0, 151.9, 140.4, 137.8, 132.7, 132.5, 132.3, 132.0, 131.9, 127.4, 127.3, 126.7, 125.0, 124.4, 109.5, 108.5, 103.5.

**HRMS (ESI^+^)** m/z: Calculated for C_23_H_15_Br_2_O_2_S [M+H]^+^: 512.9154. Found: 512.9165.

#### Scheme synthesis of Compound 21: 3-(5-((2,6-Bis(4-bromophenyl)-4*H*-pyran-4-ylidene)methyl)thiophen-2-yl)-2-cyanoacrylic acid

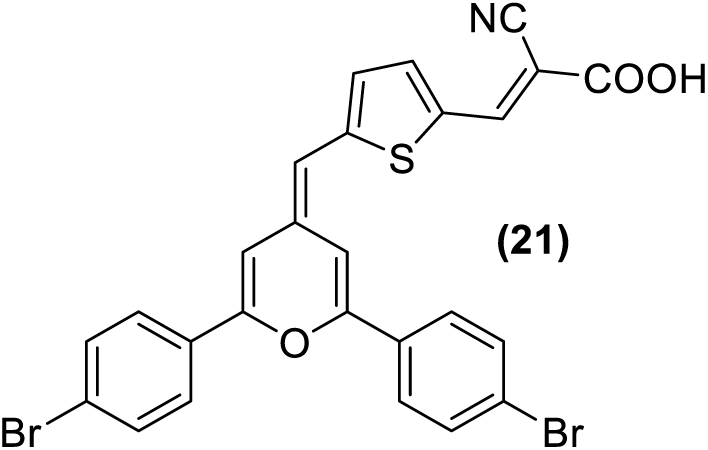

To a solution of the aldehyde prepared in the previous step (106.2 mg, 0.207 mmol) and cyanoacetic acid (35 mg, 0.411 mmol) in anhydrous acetonitrile (15 mL) under an argon atmosphere, piperidine (150 μL, 1.4 mmol) was added. The mixture was refluxed for 72 h and then cooled in an ice bath. Acetic acid (5 mL) was then added, and the resulting solid was filtered and washed with cold acetonitrile and hexane. Yield: 109.3 mg, 91%, as a dark purple solid.

**M.p.:** 220 – 222 ⁰C. **IR** (KBr) cm^-1^: 2209 C≡N), 1650 (C=O), 1540 (C=C).

**^1^H-NMR** (400 MHz, THF) δ (ppm): 8.30 (s, 1H), 7.87 (d, J = 8.7 Hz, 2H), 7.80 (d, J = 8.7 Hz, 2H), 7.74 (m, 1H), 7.66 (t, J = 8.3 Hz, 4H), 7.48 (d, J = 1.6 Hz, 1H), 7.11 (m, 1H), 6,854 (s, 1H), 6.33 (s, 1H).

**^13^C-NMR** (100 MHz, THF) δ (ppm): 164.7, 155.1, 152.8, 152.7, 146.1, 139.6, 134.2, 133.2, 133.0, 132.7, 132.6, 127.9, 127.5, 125.4, 124.7, 117.8, 110.3, 109.3, 104.3, 97.4.

**HRMS** (ESI^-^) m/z: Calculated for C_25_H_14_Br_2_NOS [M-CO_2_-H]^-^: 533.9168. Found: 533.9172. Calculated for C_52_H_29_Br_4_N_2_O_6_S_2_ [2M-H]^-^: 1156.8206. Found: 1156.8179.

#### Scheme synthesis of Compound 26

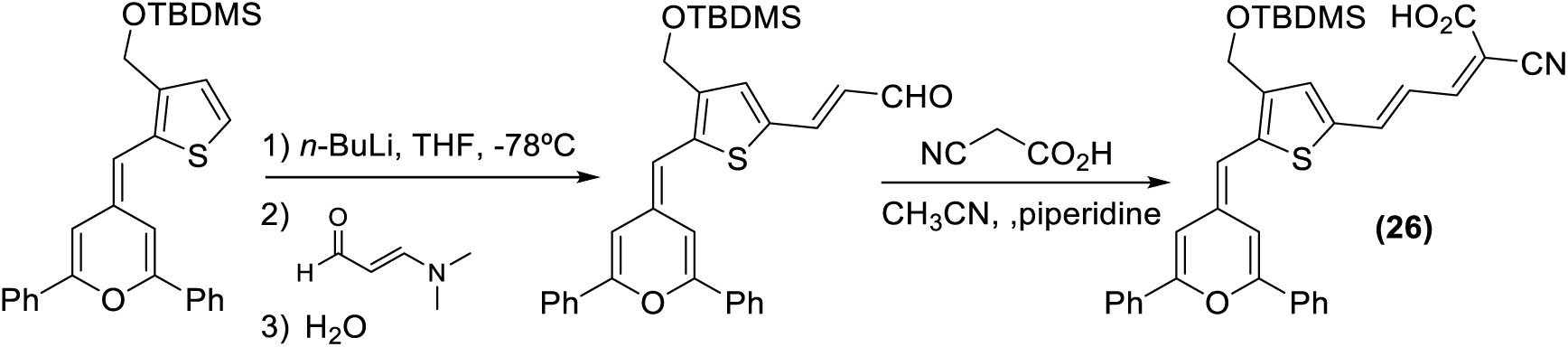

#### *tert*-butyl((2-((2,6-diphenyl-4*H*-pyran-4-ylidene)methyl)thiophen-3-yl)methoxy) dimethylsilane

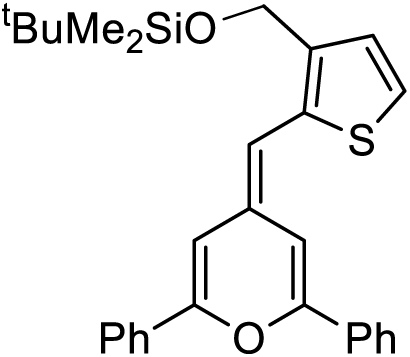

A solution of (2,6-diphenyl-4*H*-pyran-4-yl)diphenylphosphine oxide **A**^28^ (680 mg, 1.56 mmol) in anhydrous THF (12 mL) was cooled to −78°C under an Ar atmosphere. Subsequently, *n*-BuLi (1.6 M in hexanes, 1.2 mL, 1.92 mmol) was added, and the solution was stirred for 20 min at −78 °C. Then, a solution of 3-*tert*-butyldimethylsilyloxymethyl-2-thiophenecarboxaldehyde^33^ (472 mg, 1.84 mmol) in anhydrous THF (5 mL) was added, allowing the temperature to gradually return to 0 °C in 3 h. Then, a saturated solution of NH_4_Cl (15 mL) was added and the organic phase was extracted with ethyl acetate, dried over MgSO_4_ and evaporated under reduced pressure. The product was purified by silica gel column chromatography (3% ethyl acetate in hexanes). Yield: 676 mg, 91%, as a brown solid.

**M.p.:** 110−114 °C. **IR** (KBr): *cm^-^*^1^ 1652 (C=C).

**^1^H NMR** (400 MHz, CDCl_3_): *δ* 7.74−7.86 (m, 4H), 7.36−7.51 (m, 6H), 7.15 (dd, *J*=2.0 Hz, 1H), 7.13 (d, *J*=5.2 Hz, 1H), 7.08 (d, *J*=5.2 Hz, 1H), 6.44 (d, *J*=2.0 Hz, 1H), 6.07 (s, 1H), 4.76 (s, 2H), 0.94 (s, 9H), 0.10 (s, 6H).

**^13^C NMR** (100 MHz, CDCl_3_): *δ* 152.9, 150.6, 138.0, 135.7, 133.4, 133.2, 129.3, 129.0, 128.6, 128.2, 124.9, 124.4, 121.2, 108.5, 105.5, 102.6, 59.9, 26.0, ‒5.2.

**HRMS (ESI^+^):** *m/z* Calculated for [C_29_H_33_O_2_SSi]^+^: 473.1965, found: 473.1947. Calculated for [C_29_H_32_NaO_2_SSi]^+^: 495.1784, found: 495.1774. Calculated for [C_29_H_32_KO_2_SSi]^+^: 511.1524, found: 511.1516.

#### 3-(4-(((tert-butyldimethylsilyl)oxy)methyl)-5-((2,6-diphenyl-4*H*-pyran-4-ylidene)-methyl)thiophen-2-yl)acrylaldehyde

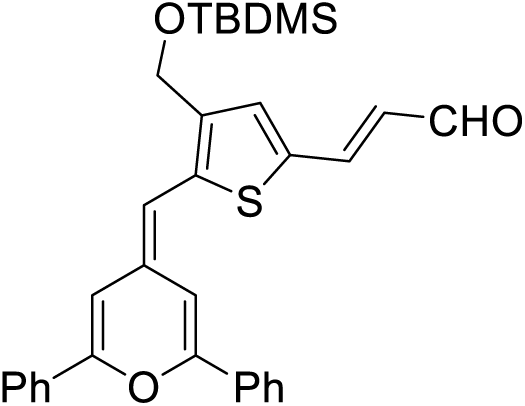

To a solution of the compound prepared in the previous step (1.04 g; 2,21 mmol) in anhydrous THF (5 mL) cooled to ‒78 °C under an argon atmosphere, *n*-BuLi (1.65 mL, 2.64 mmol) was added. The resulting mixture was stirred for 1 h to ‒78 °C and then 1 h at 0 °C. The reaction was cooled again to ‒78 °C and a solution of *N,N*-dimethylacrolein (0.66 mL, 5.96 mmol) was added. The mixture was stirred, and the temperature was allowed to slowly rise to room temperature over 3h. Then, H_2_O (0.5 mL) was added, and the reaction was stirred overnight. The THF was evaporated, and the organic phase was extracted with ethyl acetate (3 × 50 mL), washed with saturated NaCl, and dried over MgSO_4_. The solvent was evaporated under reduced pressure and the crude product was purified by silica gel column chromatography (5% ethyl acetate in hexanes). Yield: 590 mg, 51% of a red solid.

**IR** (KBr): *cm^-^*^1^ 1656 (C=O), 1114 (C−O).

**^1^H NMR** (400 MHz, CDCl_3_): δ (*ppm*) 9.60 (d, *J*=7.8 Hz, 1H), 7.88−7.73 (m, 8H), 7.58−7.40 (m, 4H), 7.21 (d, *J*=1.9 Hz, 1H), 6.50 (d, *J*=1.9 Hz, 1H), 6.46 (dd, J_1_=15.3 Hz, J_2_ = 7.8 Hz, 1H), 6.04 (s, 1H), 4.72 (s, 2H), 0.96 (s, 9H), 0.13 (s, 6H).

**^13^C NMR** (100 MHz, CDCl_3_): δ *(ppm)* 192.7, 154.7, 152.1, 144.5, 142.0, 139.5, 134.2, 133.8, 132.8, 132.7, 131.1, 130.0, 129.5, 128.9, 128.7, 125.1, 125.0, 124.6, 109.0, 105.0, 102.9, 59.7, 25.9, 18.4, ‒5.2

**HRMS (ESI^+^):** *m/z* Calculated for C_32_H_35_O_3_SSi [M+H]^+^ 527.2071; found: 527.2054.

#### Scheme synthesis of Compound 26: 5-{4-[(tert-Butyldimethylsilyloxy)methyl]-5-[(2,6-diphenyl-4*H*-pyran-4-ylidene)methyl]thiophen-2-yl}-2-cyanopenta-2,4-dienoic acid

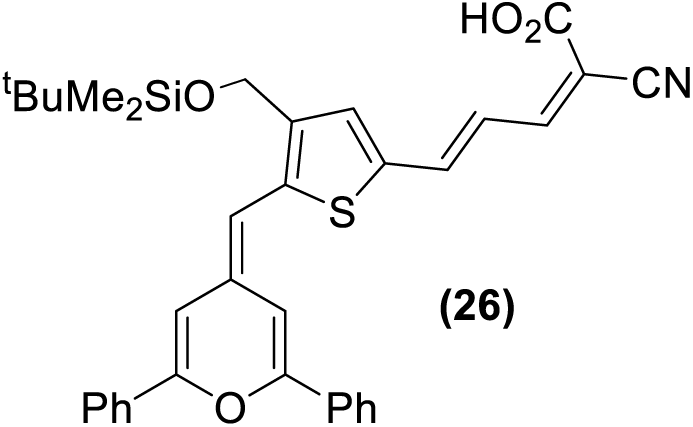

To a solution of the aldehyde prepared in the previous step (130 mg, 0.25 mmol) and cyanoacetic acid (33 mg, 0.39 mmol) in chloroform (5 mL) under an argon atmosphere, piperidine (162 μL, 1.62 mmol) was added. The mixture was refluxed for 2 h and then cooled in an ice bath. Then, acetic acid (5 mL) and H_2_O (20 mL) were added, and the organic phase was separated, dried over MgSO_4,_ and the solvent evaporated under reduced pressure. The resulting solid was washed five times with a mixture of CH_2_Cl_2_/Hexane (6:96). Yield: 142 mg, 96%, as a brown solid.

**IR (KBr):** *cm^-^*^1^ 3700−3000 (COO−H), 2216 (C≡N), 1675 (C=O), 1648 (C=C), 1234 (C-O).

**^1^H NMR** (400 MHz, DMSO-d6): *δ* 7.94−7.80 (m, 5H), 7.6 (d, *J*=14.7 Hz, 1H), 7.57−7.45 (m, 6H), 7.37 (s, 1H), 7.16 (d, J = 1.0 Hz, 1H), 6.96 (d, J=1.0 Hz, 1H), 6.77 (dd, J_1_=14.7 Hz, J_2_ =11.9 Hz, 1H), 6.28 (s, 1H), 4.70 (s, 2H), 0.89 (s, 9H), 0.09 (s, 6H).

**^13^C NMR** (100 MHz, CDCl_3_): *δ* 154.1, 151.1, 140.1, 135.6, 133.2, 133.1, 132.3, 132.1, 130.6, 130.0, 129.9, 129.3, 129.2, 125.2, 124.6, 120.9, 117.4, 109.2, 106.1, 102.6, 59.1, 26.0, 18.2, –5.0.

**EM (MALDI^+^):** *m/z* 593 C_35_H_35_NO_4_SSi [M]^+·^. HRMS (ESI^+^): *m/z* Calculated for C_35_H_36_NO_4_SSi [M+H]^+^ 594.061, found 594.2134.

#### Synthesis scheme of Compound 27: 3-(4-(((tert-butyldimethylsilyl)oxy)methyl)-5-((2,6-di-*tert*-butyl-4*H*-pyran-4-ylidene)methyl)thiophen-2-yl)-2-cyanoacrylic acid

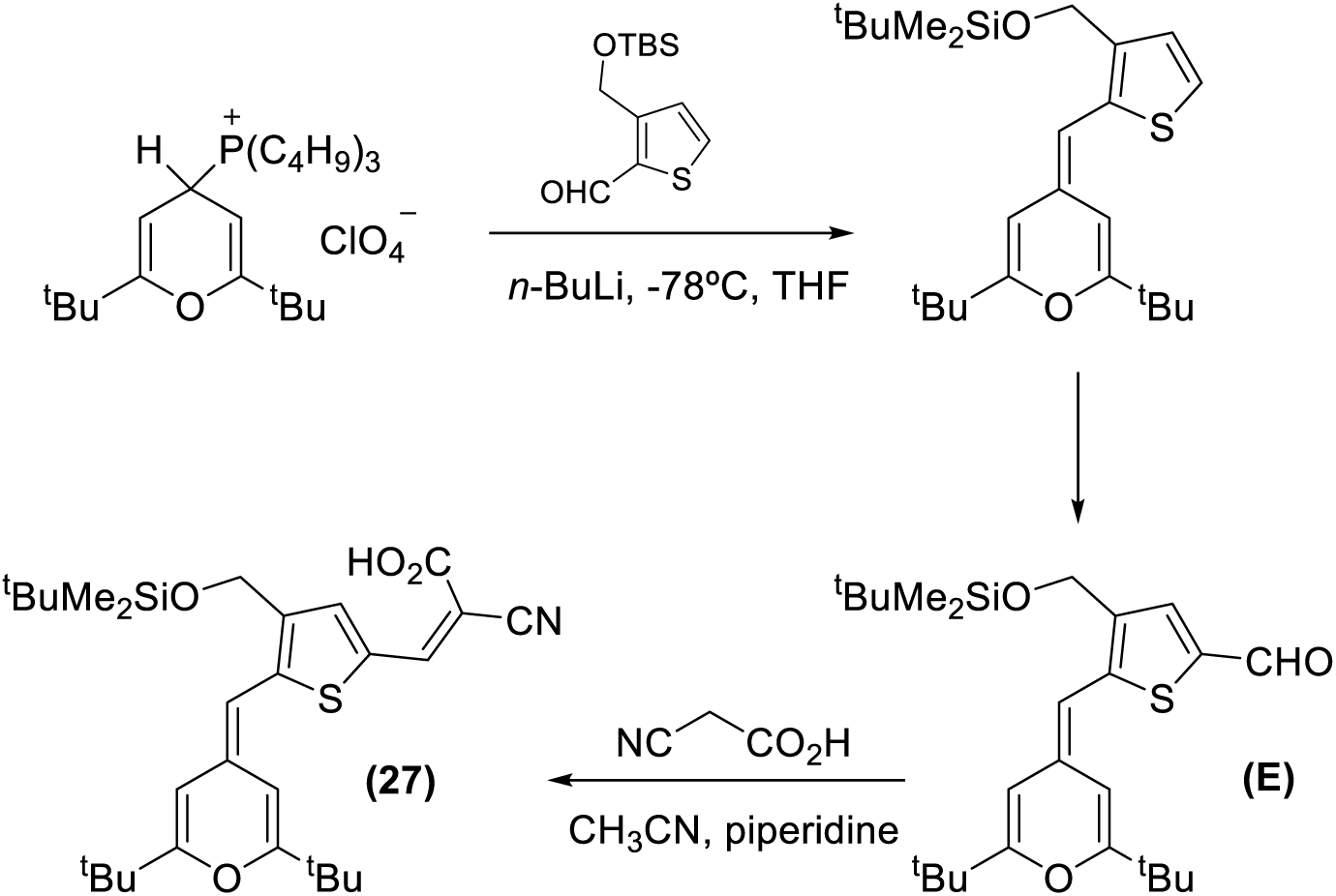

Step 1:

A solution of (2,6-di-*tert*-butyl-4*H*-pyran-4-yl)tributylphosphonium perchlorate^34^ (400 mg; 0.808 mmol) in anhydrous THF (10 mL) was prepared, purged with argon, and cooled to ‒78 °C. To the solution, *n*-BuLi (1.6 M in hexanes) (0.55 mL; 0.888 mmol) was added dropwise and the resulting mixture was stirred for 15 minutes. Then 3-(*tert*-butyldimethylsilyloxymethyl)thiophene-2-carbaldehyde^33^ (159 mg, 0.621 mmol) in anhydrous THF (5 mL) was added dropwise and the mixture was progressively heated to reach room temperature after 15 hours (TLC monitoring with 30% ethyl acetate in hexanes). A saturated solution of NH_4_Cl (20 mL) was then added to quench the reaction and the mixture was stirred for 5 minutes at room temperature. The organic layer was dried over MgSO_4,_ and the solvent was evaporated under reduced pressure. The crude product was filtered through a short silica gel column chromatography (30% ethyl acetate in hexanes) and used in the next step without additional purification. Yield: yellow oil (240 mg, 89%).

Step 2: Synthesis of compound **E**

A solution of the compound prepared in step 1 (240 mg, 0.55 mmol) in anhydrous THF (10 mL), was prepared, purged with argon and cooled to ‒45 °C. To the solution, n-BuLi (1.6 M in hexanes) (0.63 mL, 1.1 mmol) was added dropwise, and the resulting mixture was stirred for 1h. Then DMF was added (128 μl, 1.65 mmol) and the mixture was progressively heated to reach room temperature in 12h. A saturated solution of NH_4_Cl (20 mL) was added and the mixture was stirred for 5 minutes. The solution was extracted with CH_2_Cl_2_ (2 × 30 mL), dried over MgSO_4_ and the solvent was evaporated under reduced pressure. Pure compound was obtained by silica gel column chromatography (10% ethyl acetate in hexanes). Yield: 174 mg, 68%) as orange oil.

**^1^H NMR** (400 MHz, CDCl_3_) δ 9.75 (s, 1H), 7.65 (s, 1H), 6.55 (d, J = 1.6 Hz, 1H), 5.81-5.78 (m, 2H), 4.69 (s, 2H), 1.27 (s, 9H), 1.22 (s, 9H), 0.94 (s, 9H), 0.11 (s, 6H)

**^13^C NMR** (100 MHz, CDCl_3_) δ 181.9, 167.3, 164.2, 147.6, 137.9, 137.8, 136.1, 134.4, 105.6, 101.9, 99.8, 59.8, 36.0, 35.5, 27.9, 27.8, 25.9, 18.4, ‒5.2.

Step 3: Synthesis of compound **27**

To a solution of the aldehyde prepared in step 2 (167 mg, 0.362 mmol) and 2-cyanoacetic acid (48 mg; 0.565 mmol) in chlorofom (30 mL) was added piperidine (243 µL; 2.4 mmol). The mixture was refluxed for 20 hours under argon atmosphere and then cooled down to room temperature. The solvent was then evaporated under reduced pressure. Pure compound was obtained by reverse C18 column chromatography (CH_3_CN/NH_4_AcO (20 mM), 1/1). Before evaporation of the solvent, a drop of diluted AcOH was added in all fractions. Yield: 120 mg, 62%, purple solid.

**^1^H NMR** (400 MHz, CDCl_3_) δ 8.20 (s, 1H), 7.65 (s, 1H), 6.75 (brs, 1H), 5.89 (m, 2H), 4.70 (s, 2H), 1.31 (s, 9H), 1.24 (s, 9H), 0.94 (s, 9H), 0.11 (s, 6H)

**^13^C NMR** (100 MHz, CDCl_3_) δ 169.8, 168.8, 165.5, 150.9, 146.5, 140.2, 138.5, 136.5, 129.1, 117.1, 106.2, 102.5, 100.8, 90.7, 59.6, 36.2, 35.7, 27.8, 25.9, 18.3, ‒5.2.

#### Synthesis scheme of Compound 30: (3-cyano-4-(2-(5-((2,6-diphenyl-4*H*-pyran-4-ylidene)methyl)-3,4-dihexylthiophen-2-yl)vinyl)-5,5-dimethylfuran-2(5H)-ylidene) malononitrile

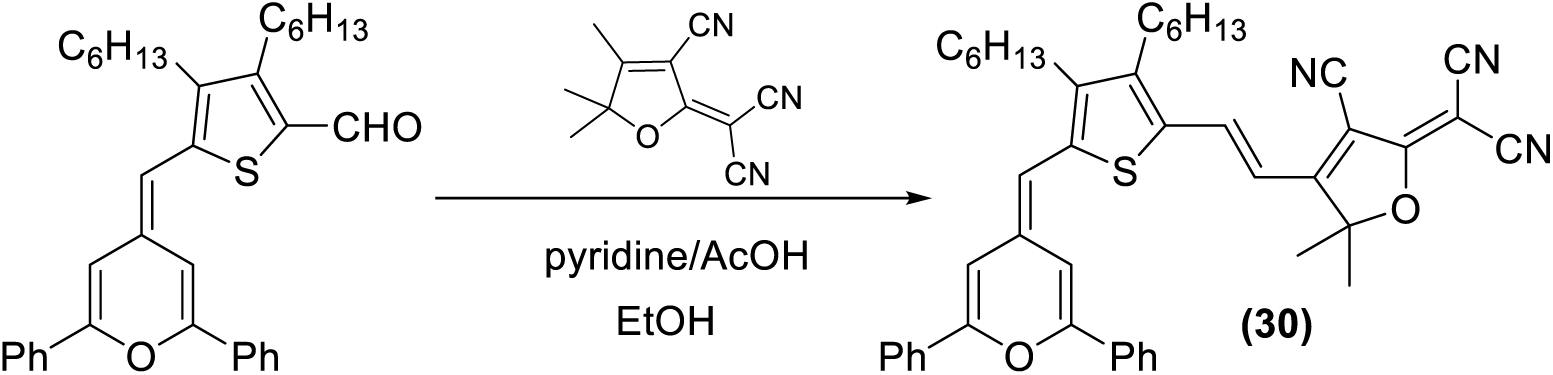

To a solution of the described aldehyde^13^ (289.2 mg, 0.55 mmol) in ethanol (10 mL) 2-(3-cyano-4,5,5-trimethylfuran-2(5*H*)-ylidene)malononitrile^35^ (100 mg, 0.55 mmol), pyridine (440 μl, 5.40 mmol) and acetic acid (217 μl, 3.78 mmol) were added and the mixture was refluxed under argon atmosphere for 24h. The reaction was stirred at 0 °C for 1h, resulting in the formation of a precipitate, which was filtered and washed with cold hexane and a mixture of hexane/CH_2_Cl_2_ (9:1). Yield: 172.3 mg, 44% of a dark red solid.

**M.p.: 210-212 °C**

**^1^H NMR** (300 MHz, CDCl_3_) δ 8.07 (d, *J* = 15.0 Hz, 1H), 7.98 – 7.76 (m, 4H), 7.64 – 7.43 (m, 6H), 7.40 (d, *J* = 2.0 Hz, 1H), 6.68 (d, *J* = 2.0 Hz, 1H), 6.41 (d, *J* = 15.0 Hz, 1H), 6.18 (s, 1H), 2.70 (t, J = 7.7 Hz, 2H), 2.62 (t, J = 8.8 Hz, 2H), 1.73 (s, 6H), 1.65 – 1.20 (m, 16H), 0.91 (d, *J* = 6.5 Hz, 6H).

**^13^C NMR** (75 MHz, CDCl_3_) *δ* 176.5, 172.3, 156.7, 154.0, 153.4, 148.7, 142.1, 136.9, 135.4, 132.3, 132.1, 132.0, 130.8, 130.2, 129.2, 129.0, 125.7, 125.0, 113.2, 112.5, 112.4, 110.1, 108.6, 106.5, 104.0, 96.1, 78.4, 31.9, 31.7, 31.6, 30.5, 29.6, 29.6, 28.0, 27.2, 26.8, 22.7, 22.6, 14.1, 14.1.

**HRMS (ESI^+^):** m/z calculated for C_46_H_48_N_3_O_2_S [M+H]^+·^ 706.3462; found 706.3413

#### Synthesis scheme of Compound 34: 3-(2,6-diphenyl-4*H*-pyran-4-ylidene)-2-phenylprop-1-ene-1,1,3-tricarbonitrile

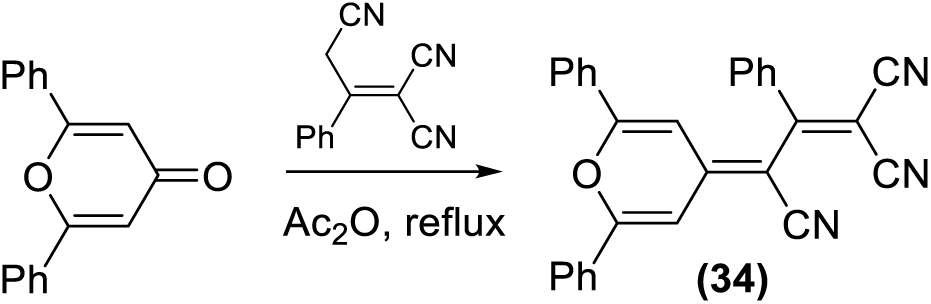

To a solution of 2,6-diphenyl-4H-pyran-4-one^36^ (300 mg, 1.208 mmol) in acetic anhydride (1.50 mL) was added 2-phenylprop-1-ene-1,1,3-tricarbonitrile^37^ (235 mg, 1.21 mmol) and the mixture was refluxed under an argon atmosphere for 1h 30min. The reaction was cooled to room temperature and then to 0 °C for 1h. The precipitate was filtered a washed three times with cold ether. Yield: 357 mg, 70%, light pink solid.

**^1^H NMR** (400 MHz, CDCl_3_, 45 °C) δ (ppm): 7.99-7.85 (m, 2H), 7.64-7.43 (m, 14H), 6.53 (d, J = 2.0 Hz, 1H)

**^13^C NMR** (100 MHz, CDCl_3_, 45 °C) δ 167.3, 161.4, 160.2, 152.5, 135.1, 133.0, 132.5, 132.4, 130.8, 130.7, 130.4, 129.8, 129.7, 129.6, 126.4, 126.4, 117.3, 114.6, 113.8, 88.4, 79.0

#### Synthesis scheme of Compound 35: 2-phenyl-3-(1,2,6-trimethylpyridin-4(1*H*)-ylidene)prop-1-ene-1,1,3-tricarbonitrile

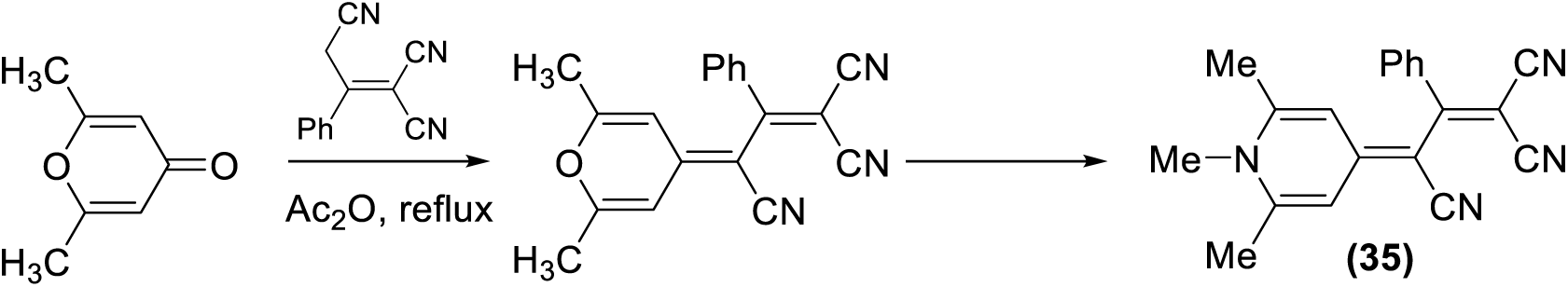

Step 1:

To a solution of 2,6-dimethyl-4H-pyran-4-one (commercially available) (117 mg, 0.941 mmol) in acetic anhydride (1.16 mL) was added 2-phenylprop-1-ene-1,1,3-tricarbonitrile^37^ (181 mg, 0.937 mmol) and the mixture was refluxed under argon atmosphere for 1h. The crude of the reaction was extracted with a mixture of CH_2_Cl_2_/H_2_O and the organic phase washed with H_3_PO_4_ (2%, 30 mL), NaOH (10%, 30 mL), saturated NaCl (30 mL) and the solvent was evaporated under reduced pressure. Yield: 265mg, 94%.

**^1^H NMR** (400 MHz, CDCl_3_, 45 °C) δ (ppm): 7.62-7.46 (m, 5H), 6.79 (d, J = 2.2 Hz, 1H), 5.96 (d, J = 2.2 Hz, 1H), 2.36 (s, 3H), 2.25 (s, 3H)

Step 2:

To a solution of the product prepared in step 1 (260 mg, 0.869 mmol) in ethanol (2 mL) a solution of dimethylamine (870 μl of a 40% solution in H_2_O) was added. The mixture of reaction was refluxed for 45 min (the solution is dark red) and then cooled to 0 °C. The red precipitate was filtered, washed with 0.5 mL of cold ethanol and dried. Yield: 111 mg, 41% of a red solid.

**^1^H NMR** (400 MHz, CDCl_3_) δ (ppm): 7.57-7.42 (m, 5H), 6.91 (s, 2H), 3.72 (s, 3H), 2.51 (s, 6H)

**^13^C NMR** (100 MHz, CDCl_3_) δ (ppm) δ 167.2, 152.8, 149.3, 136.4, 131.6, 130.4, 129.0, 120.3, 120.1, 117.9, 116.9, 80.1, 62.6, 56.8, 37.2, 21.6.

**MS (MALDI):** m/z 313.149 [M+H] ^+^

#### Synthesis scheme of Compound 37, Compound 38 and Compound 39

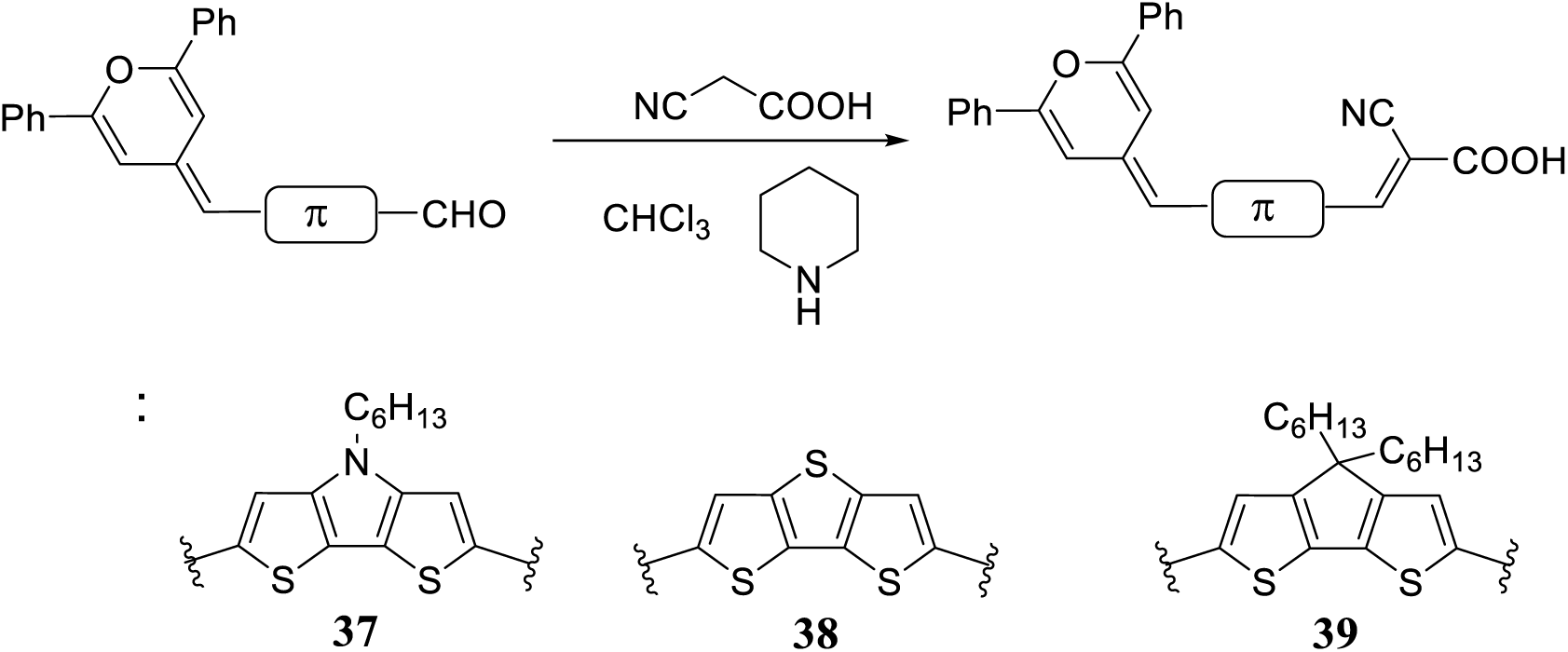

#### Compound 37: 2-cyano-3-(6-((2,6-diphenyl-4*H*-pyran-4-ylidene)methyl)-4-hexyl-4*H*-dithieno[3,2-*b*:2’,3’-*d*]pyrrol-2-yl)acrylic acid

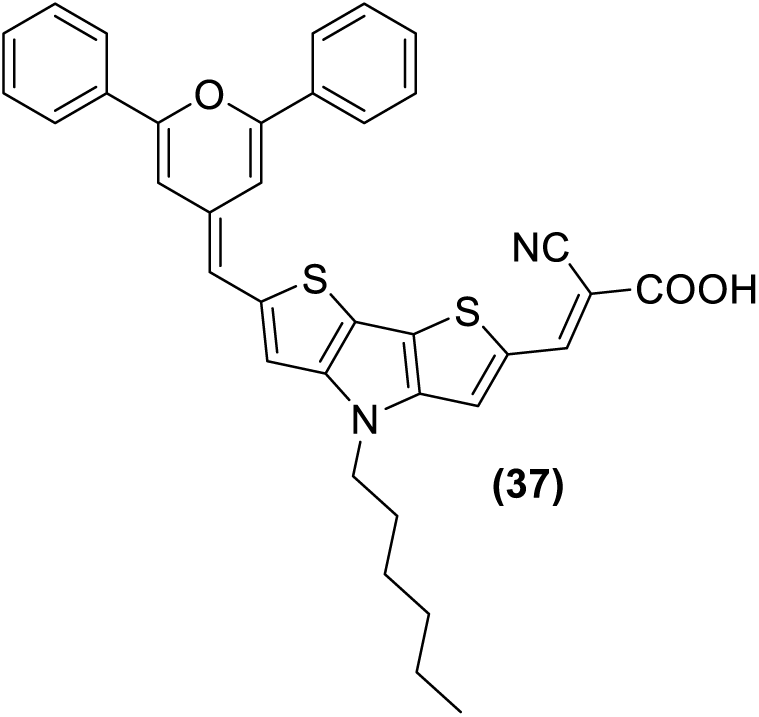

To a solution of *N*-hexyl-6-(2,6-diphenyl-4*H*-pyran-4-ylidenemethyl)dithieno[3,2-*b*:2’,3’-*d*]pyrrole-2-carbaldehyde^38^ (0.069 g, 0.130 mmol) and cyanoacetic acid (0.016 g, 0.19 mmol) in chloroform (4 mL) under an argon atmosphere, piperidine (85 μL, 0.85 mmol) was added. The mixture was refluxed for 48 h and then cooled in an ice bath. The resulting solid was filtered, washed with cold hexane, and subsequently washed with an aqueous solution of 0.1 N HCl and water. The product was obtained as a dark purple solid (0.058 g, 0.096 mmol, 75%).

**M.p.:** (°C): 174–176. **IR** (KBr): ρ̅ (cm^-1^) 2209 (C≡N), 1652 (C=O), 1570 (C=C).

**^1^H NMR** (500 MHz, DMSO-*d*_6_, 40 °C, presat. water) δ (ppm): 8.39 (s, 1H), 8.06 (s, 1H), 8.01–7.82 (m, 4H), 7.69–7.41 (m, 6H), 7.28 (s, 1H), 7.24 (s, 1H), 6.92 (s, 1H), 6.34 (s, 1H), 4.38–4.24 (m, 2H), 1.91–1.74 (m, 2H), 1.36–1.15 (m, 6H), 0.87–0.74 (m, 3H).

**^13^C NMR**: not registered due to the poor solubility of the product.

**HRMS (ESI^+^):** m/z calculated for C_36_H_30_N_2_O_3_S_3_ [M^+·^] 602.1692; found 602.1640.

#### Compound 38: 2-cyano-3-(6-((2,6-diphenyl-4*H*-pyran-4-ylidene)methyl)dithieno-[3,2-*b*:2’,3’-*d*]thiophen-2-yl)acrylic acid

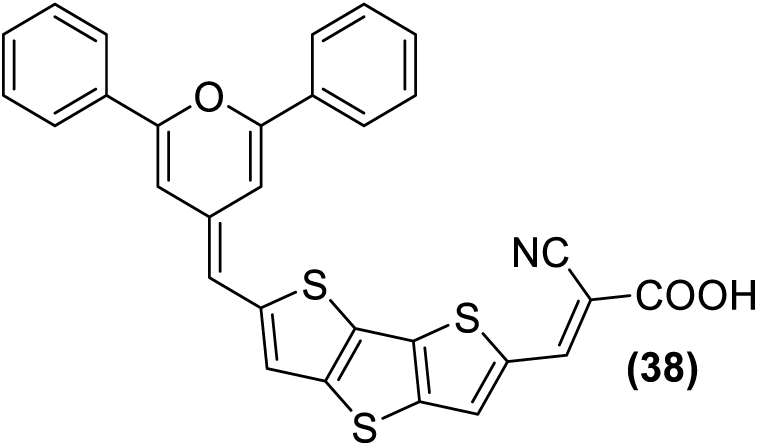

To a solution of 6-(2,6-diphenyl-4*H*-pyran-4-ylidenemethyl)dithieno[3,2-*b*:2’,3’-*d*]thiophene-2-carbaldehyde^39^ (0.050 g, 0.14 mmol) and cyanoacetic acid (0.014 g, 0.17 mmol) in chloroform (3.5 mL) under an argon atmosphere, piperidine (80 μL, 0.80 mmol) was added. The mixture was refluxed for 48h, then cooled in an ice bath. The resulting solid was collected by filtration, washed successively with a cold mixture of hexane/dichloromethane (8:2), an aqueous solution of 0.1 N HCl, and water. The product was obtained as a purple solid (0.032 g, 0.060 mmol, 56%).

**M.p.:** (°C): 215–217. **IR** (KBr): ρ̅ (cm^-1^) 2213 (C≡N), 1652 (C=O), 1570 (C=C).

**^1^H NMR** (500 MHz, DMSO-*d*_6_, 77°C, presat. water) δ (ppm): 8.51 (s, 1H), 8.33 (s, 1H), 8.03–7.95 (m, 2H), 7.93–7.85 (m, 2H), 7.64–7.45 (m, 7H), 7.18 (s, 1H), 6.92 (s, 1H), 6.34 (s, 1H).

**^13^C NMR**: not registered due to the poor solubility of the product.

**HRMS (MALDI^+^):** m/z calculated for C_30_H_17_NO_3_S_3_ [M^+·^] 535.0365; found 535.0439.

#### Compound 39: 2-cyano-3-(6-((2,6-diphenyl-4*H*-pyran-4-ylidene)methyl)-4,4-dihexyl-4*H*cyclopenta[2,1-*b*:3,4-*b*’]dithiophen-2-yl)acrylic acid

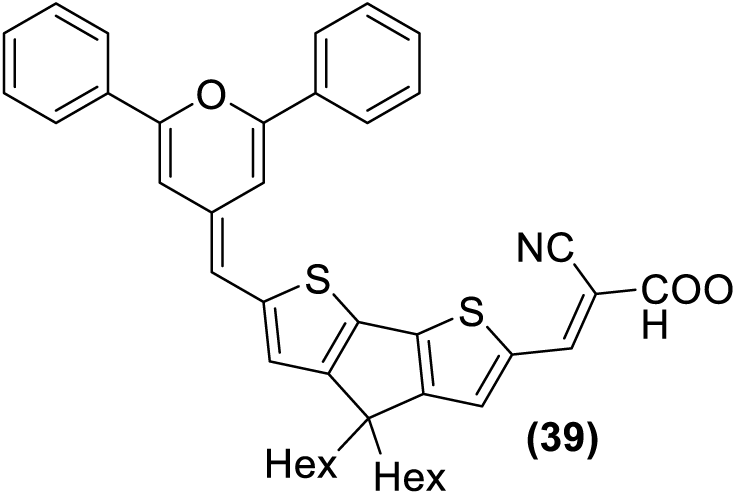

To a solution of 4,4-dihexyl-6-(2,6-diphenyl-4*H*-pyran-4-ylidenemethyl)-cyclopenta[2,1-*b*:3,4-*b*’]dithiophene-2-carbaldehyde^38^ (0.061 g, 0.098 mmol) and cyanoacetic acid (0.013 g, 0.15 mmol) in chloroform (3 mL) under argon atmosphere, piperidine (65 μL, 0.65 mmol) was added. The mixture was refluxed for 87 h and then cooled in an ice bath. The resulting solid was filtered and washed with hexane, followed by an aqueous solution of 0.1 N HCl and water. The product was obtained as a dark blue solid (0.034 g, 0.049 mmol, 50%).

**M.p.**: (°C): 128–130. IR (KBr): ρ̅ (cm^-1^) 2209 (C≡N), 1654 (C=O), 1570 (C=C).

**^1^H NMR** (400 MHz, CD_2_Cl_2_) δ (ppm): 8.28 (s, 1H), 7.95–7.89 (m, 2H), 7.84–7.78 (m, 2H), 7.61–7.34 (m, 7H), 7.22 (d, *J* = 1.4 Hz, 1H), 6.86 (s, 1H), 6.54 (d, *J* = 1.4 Hz, 1H), 6.20 (s, 1H), 1.97–1.81 (m, 4H), 1.23–1.08 (m, 12H), 1.03–0.92 (m, 4H), 0.85–0.77 (m, 6H).

**^13^C NMR**: not registered due to the poor solubility of the product.

**HRMS (ESI^+^):** m/z calculated for C_43_H_43_NO_3_S_2_ [M^+·^] 685.2684; found 685.2697; calculated for C_43_H_44_NO_3_S_2_ [M+H]^+^: 686.2757; found 686.2689.

#### Scheme synthesis of Compound 42: 2-cyano-4-(2,6-di-tert-butyl-4*H*-pyran-4-ylidene)but-2-enoic acid

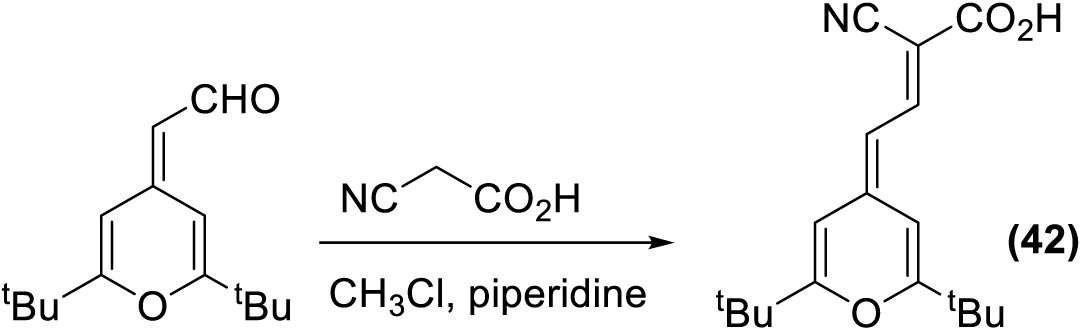

To a solution of 2-(2,6-di-tert-butyl-4*H*-pyran-4-ylidene)acetaldehyde^30^ (112 mg, 0.48 mmol) and cyanoacetic acid (63.6 mg, 0.748 mmol) in chloroform (15 mL) under argon atmosphere, piperidine (317μl, 3.18 mmol). The mixture was refluxed for 20h and the solvent evaporated under reduced pressure. Pure compound was obtained by reverse C18 column chromatography (MeOH/NH_4_AcO (20 mM), 3/7). Before evaporation of the solvent, a drop of 10% AcOH was added in all fractions. Yield: 116 mg, 80%, orange solid.

**M.p:** 203-204 °C

**^1^H NMR:** (400 MHz, CDCl_3_) δ 1.27 (s, 18H), 1.30 (s, 18H), 6,0 (d, *J* = 6 Hz, 1H), 6,3 (dd, *J* = 6Hz, *J* = 6.3Hz, 2H), 8.25 (d, *J* = 8.3Hz, 1H)

**^13^C NMR:** (100 MHz, THF) δ 25.2, 25.3, 98.4, 104.6, 105.8, 146.9

**HRMS (ESI^+^)** m/z: Calculated for C_18_H_23_O_3_N: 301.1672 found: 301.1152

#### Synthesis scheme of Compound 46: 3-(4-(((*tert*-butyldiphenylsilyl)oxy)methyl)-5-((2,6-di-tert-butyl-4*H*-pyran-4-ylidene)methyl)thiophen-2-yl)-2-cyanoacrylic acid

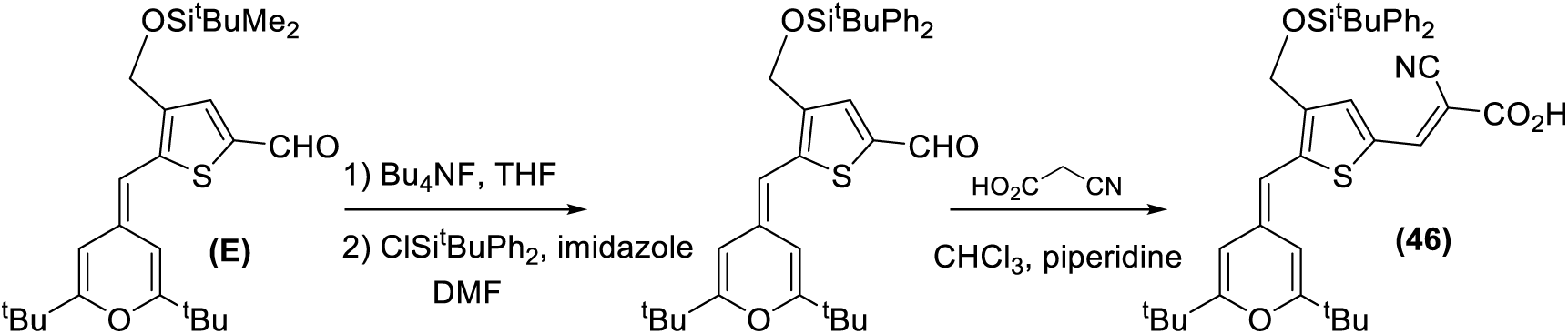

Step 1: To a solution of aldehyde **E** (see above synthesis of compound 27) (188 mg, 0.41 mmol) in anhydrous THF (10 mL) was added Bu_4_NF (1.0 M in THF) (0.815 mL, 0.815 mmol). The mixture was stirred at 0 °C under an argon atmosphere for 1h. Subsequently, the solvent was evaporated under reduced pressure, and the resulting crude product was used directly in step 2 without further purification.

The crude was dissolved in anhydrous DMF (8 mL) and imidazole (140.7 mg, 0.857 mmol) and ClSi^t^BuPh_2_ (0.129 mL, 0.49 mmol) were added. The mixture was stirred at room temperature overnight and then 30 mL of water was added. The solution was extracted with ethyl acetate (3 × 15 mL), washed with saturated NaCl and the organic phase dried over MgSO_4_. The pure compound was obtained by silica gel column chromatography (10% ethyl acetate in hexanes). Yield: 141 mg, 60%).

**^1^H NMR** (400 MHz, CD_2_Cl_2_) δ 9.71 (s, 1H), 7.75-7.62 (m, 4H), 7.60 (s, 1H), 7.51-7.34 (m, 6H), 6.51 (d, *J* = 1.9 Hz, 1H), 5.72-5.70 (m, 2H), 4.73 (s, 2H), 1.27 (s, 9H), 1.20 (s, 9H), 1.07 (s, 9H)

**^13^C NMR** (100 MHz, CD_2_Cl_2_) δ 182.0, 167.7, 164.6, 148.0, 138.3, 137.6, 136.6, 136.0, 134.8, 133.7, 130.2, 128.2, 105.9, 102.2, 100.1, 36.3, 35.8, 28.0, 27.9, 27.0, 19.5

Step 2: To a solution of the aldehyde obtained in step 1 (106 mg, 0.181 mmol) and cyanoacetic acid (24.2 mg, 0.284 mmol) in chloroform (15 mL) under an argon atmosphere, piperidine (132 μl, 1.20 mmol). The reaction mixture was refluxed for 20h, after which the solvent was removed under reduced pressure. The pure compound was isolated by reverse C18 column chromatography (MeOH/NH_4_AcO (20 mM), 1/1). Before solvent removal, a drop of 10% AcOH was added in all collected fractions. Yield: 71.5 mg, 60% as a purple solid.

**^1^H NMR** (400 MHz, TFH-d8) δ 8.21 (s, 1H), 7.76-7.69 (m, 5H), 7.44-7.37 (m, 6H), 6.74 (bs, 1H), 5.88-5.84 (m, 2H), 4.78 (s, 2H), 1.30 (s, 9H), 1.22 (s, 9H), 1.08 (s, 9H)

**^13^C NMR** (100 MHz, THF-d8) δ 168.5, 165.2, 165.1, 148.8, 146.0, 140.0, 138.6, 138.3, 136.6, 135.4, 134.3, 131.0, 130.8, 128.8, 126.1, 117.7, 107.0, 103.7, 101.3, 95.7, 61.3, 37.0, 36.4, 35.3, 28.3, 28.3, 27.4.

#### Scheme synthesis of Compound 50 5-(4-diethylamino)benzylidene-2-oxo-4-phenyl-2,5-dihydrofuran-3-carbonitrile ( see Table S1)

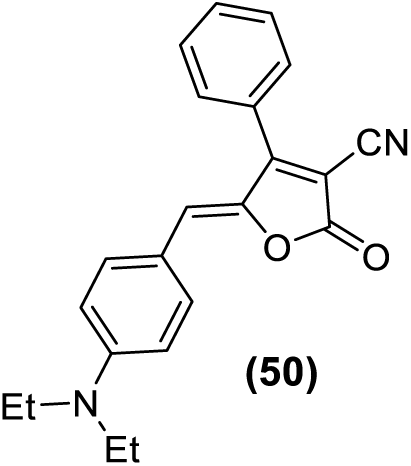

A mixture of 270 mg (1,5 mmol) of 4-diethylaminobenzaldehyde (commercially available) and 4-phenyl-2-oxo-2,5-dihydrofuran-3-carbonitrile^40^ (278 mg, 1.5 mmol) in absolute ethanol (8 mL) was refluxed under an argon atmosphere and protected from light for 14h. After cooling to room temperature, the resulting solid was collected by filtration, washed with cold ethanol and dried, to yield a dark violet solid (380 mg, 74%).

**M.p.:** (°C): 211−217. **IR** (nujol): ρ̅ (cm^-1^) 2214 (C≡N), 1758 (C=O), 1593 (C=C), 1544 and 1528 (C=C, Ar.).

**^1^H NMR** (400 MHz, CDCl_3_) δ (ppm): 7.78−7.73 (m, 2H), 7.64−7.55 (m, 5H), 6.71−6.64 (m, 2H), 6,39 (s, 1H), 3,46 (q, *J* = 7.1 Hz, 4H), 1.23 (t, *J* = 7.1 Hz, 6H).

**^13^C NMR** (100 MHz, CDCl_3_) δ (ppm): 165.4, 162.1, 150.2, 142.2, 135.0, 131.5, 129.3, 128.9, 128.4, 123.6, 120.1, 112.9, 111.8, 93.8, 44.8, 12.6.

**HRMS (ESI^+^):** m/z calculated for C_22_H_21_N_2_O_2_ ([M+H]^+^) 345.1598, found 345.1604; calculated for C_22_H_20_N_2_NaO_2_ ([M+Na]^+^) 367.1417, found 367.1385

#### Scheme synthesis of Compound 52: 2-cyano-3-(5-((4,5-dimethyl-1,3-dithiol-2-ylidene)methyl) thiophen-2-yl)acrylic acid (Table S1)

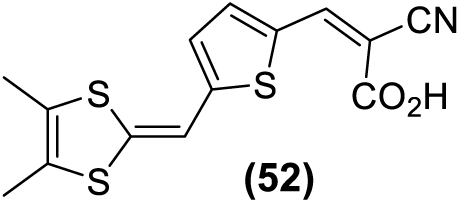

The starting aldehyde was synthesized in high yield following a previously reported procedure^41^. To a solution of aldehyde (200mg, 0,787 mmol) and 2-cyanoacetic acid (87 mg, 1.02 mmol) in anhydrous acetonitrile (20 mL), piperidine (30 mL) was added. The mixture was refluxed for 24 hours under an argon atmosphere, then cooled down to 0 ^°^C and maintained at this temperature for 30 minutes. The resulting solid was collected by filtration, washed with a cold mixture of 10% dichloromethane in hexanes (30 mL) and dried. Yield: dark violet solid (154 mg, 61%).

**M.p**: 214-215 ^°^C. IR (KBr): cm^-1^ 2216 (C=N), 1675 (C=O), 1562 (C=C).

**^1^H NMR** (400 MHz, DMSO-d6) 8 (ppm): 8.38 (s, 1H), 7.91 (d, *J* = 4.1 Hz, 1H), 7.19 (s, 1H), 7.06 (d, *J* = 4.1 Hz,1H), 2,12 (s, 3H), 2,06 (s, 3H).

**^13^C-NMR** (100 MHz, DMSO-d6) 8 (ppm): 164.2, 150.7, 145.8, 141.7, 140.9, 132.5, 124.4, 124.1, 124.0, 117.1, 105.1, 94.1, 12.6

**HRMS (ESI+)** m/z: Calculated for C_14_H_12_NO_2_S_3_: 322.0025. Found: 322.0027

## Acknowledgments

This research was funded by a fellowship from the Spanish Government (Programa de Formación de Profesorado Universitario) Ref. FPU18/03873 to J.M.E-A. This work was supported by internal funds (J.A.A. and S.R.-G). R.A. and S.F. were funded by Gobierno de Aragón-FEDER Fondo Social Europeo (E47_23R) and the University of Zaragoza (UZ2023-CIE-01).

The authors would like to thank Iñigo Lasa (Universidad Pública de Navarra) for providing pMAD and the protocol for genetic manipulation of *S. aureus*.

## Authors contributions

CRediT (Contributor Roles Taxonomy) has been applied for author contribution. Conceptualization: S.F., J.A.A., S.R.-G.; Investigation: J.M.E.-A., R.A.-R., A.L.; Formal Analysis: J.M.E.-A., R.A.-R., A.L.; Resources: R.A., S.F.; Supervision: S.F., J.A.A., S.R.-G.; Writing – original draft: J.M.E.-A., S.F.; Writing – review and editing: J.M.E.-A., R.A., S.F., J.A.A., S.R.-G. All authors have reviewed and agreed to the final version of the manuscript.

## Potential conflicts of interest

Authors declare no conflict of interest.

## Notes

### Competing Interest Statement

The authors have declared no competing interest.

