## Supplementary material for "Novel antimicrobial activity of photoactive compounds originally synthesized for industrial applications": Table S1

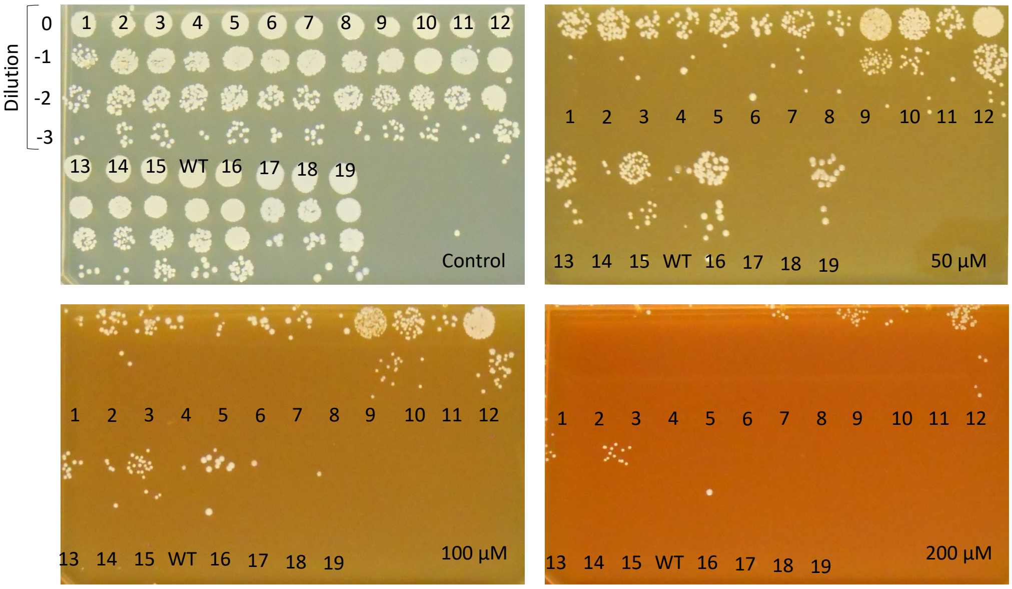


**Fig. S1. Drop dilution Compound 18 susceptibility testing of nineteen *S. aureus* mutants.** Compound18 concentrations are indicated in µM; a Compound 18-free plate was included as a control. Mutants 17 and 18 were discarded for subsequent WGS given that they behaved as the parental wild-type (WT) *S. aureus* ATCC 29213.

**Table S1. Activity of the panel of 21 compounds representing the chemical diversity of the PMM compound collection against *Mycobacterium tuberculosis*.** MICs were determined in 7H9-0.2% glycerol-ADC, bacteria were incubated for 6 days before the addition of MTT and one additional day to allow for MTT conversion to formazan.

| Compound | Reference | MIC (μM) |
|---|---|---|
| 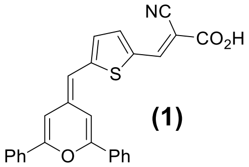 (1) | 1 | 50 |
| 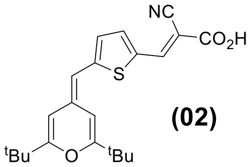 (02) | 1 | >50 |
| 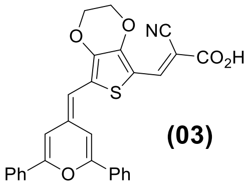 (03) | This work | >50 |
| 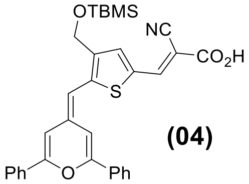 (04) | 2 | >50 |
| 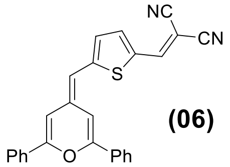 (06) | This work | >50 |
| 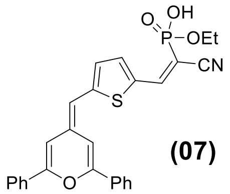 (07) | This work | >50 |
| 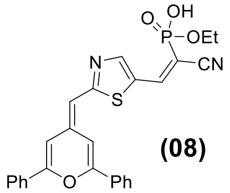 (08) | 3 | >50 |
| 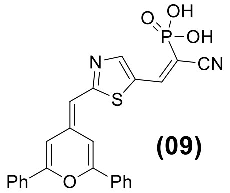 (09) | 3 | >50 |

| | | |
|---|---|---|
| 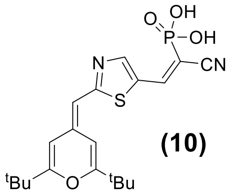 (10) | This work | >50 |
| 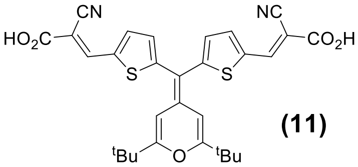 (11) | 4 | >50 |
| 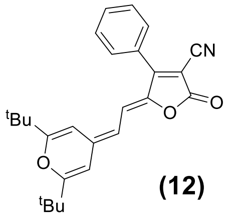 (12) | 5 | >50 |
| 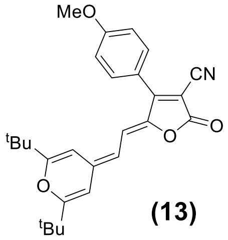 (13) | This work | >50 |
| 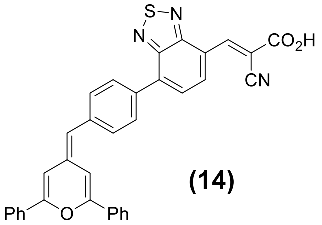 (14) | This work | >50 |
| 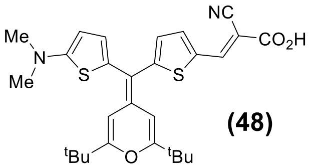 (48) | 4 | >50 |
| 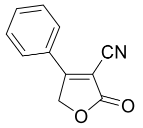 (49) | 6 | >50 |
| 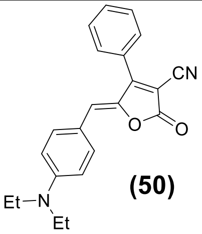 (50) | This work | >50 |
| 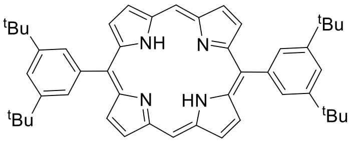 (51) | 7 | >50 |

| | | |
|---|---|---|
| 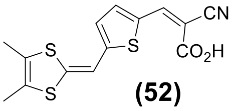 **(52)** | This work | >50 |
| 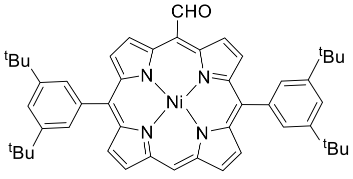 **(53)** | 8 | >50 |
| 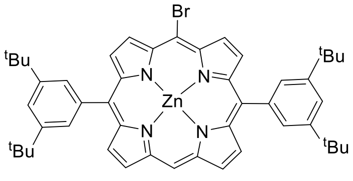 **(54)** | 9 | >50 |
| 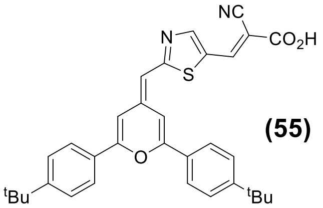 **(55)** | 10 | >50 |

**Table S3. Bacterial strains used in the screening of the Photoactive Molecular Materials compound collection.**

| Strain | Type | Incubation time | Culture medium |
|---|---|---|---|
| *Corynebacterium diphteriae* ATCC 39255 | Gram-positive | 24 h | BHI |
| *Corynebacterium glutamicum* ATCC 13032 | Gram-positive | 48 h | Müller-Hinton II |
| *Enterococcus faecalis* ATCC 19433 | Gram-positive | 24 h | Müller-Hinton II |
| *Staphylococcus aureus* ATCC 29213 (MSSA) | Gram-positive | 24 h | Müller-Hinton II |
| *Staphylococcus aureus* ATCC 43300 (MRSA) | Gram-positive | 24 h | Müller-Hinton II |
| *Staphylococcus epidermidis* CECT 231 | Gram-positive | 24 h | Müller-Hinton II |
| *Klebsiella pneumoniae* ATCC 13883 | Gram-negative | 24 h | Müller-Hinton II |
| *Pseudomonas aeruginosa* ATCC 15442 | Gram-negative | 24 h | Müller-Hinton II |
| *Salmonella typhimurium* ATCC 14028 | Gram-negative | 24 h | Müller-Hinton II |
| *Mycobacteroides abscessus subsp. abscessus* ATCC 19977 | mycobacteria | 72 h | 7H9-0.2% glycerol-ADC |
| *M. abscessus subsp. bolletii* CCUG 50184 | mycobacteria | 72 h | 7H9-0.2% glycerol-ADC |
| *M. abscessus subsp. massiliense* CCUG 48898 | mycobacteria | 72 h | 7H9-0.2% glycerol-ADC |
| *Mycobacterium avium* ATCC 25291 | mycobacteria | 72 h | 7H9-0.2% glycerol-ADC |
| *Mycolicibacterium smegmatis* $mc^2155$ | mycobacteria | 72 h | 7H9-0.2% glycerol-ADC |

**Table S4. Single shot assay of the 4H-pyran-4-ylidene derivatives.** Only derivatives active against at least one Gram-positive bacterial species are shown. Compounds active in two assay replicates are shown in black; grey squares indicate compounds were active in only one of the replicates, which were also selected for further analysis. All 4H-pyran-4-ylidene derivatives were inactive against Gram-negative bacteria and mycobacteria (**Table S2**).

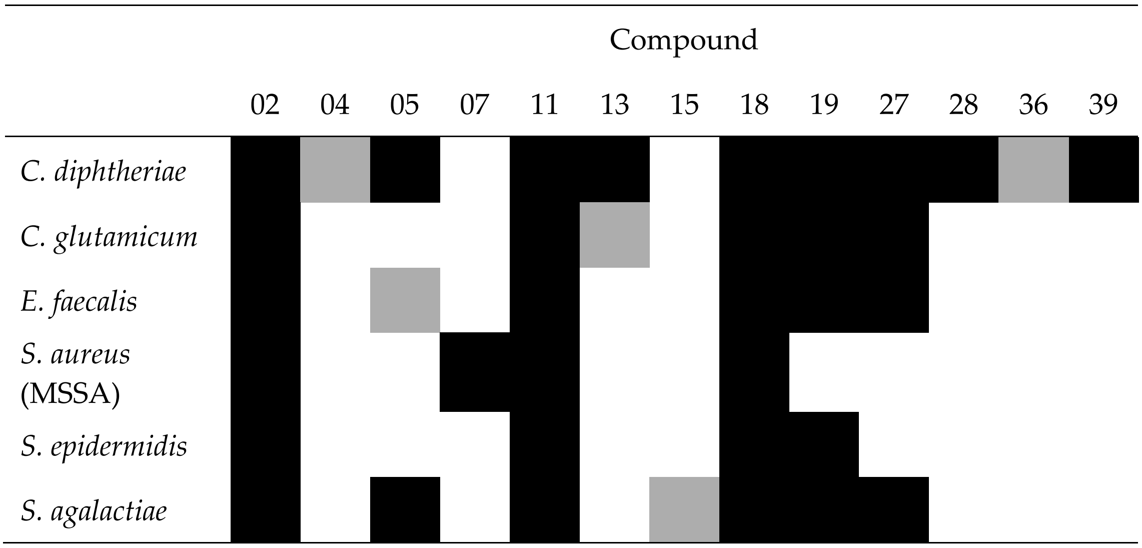

**Table S5. Mutations identified in *S. aureus* resistant mutants.** Mutants 1-15 were isolated from liquid cultures. Mutants 16 and 19 were isolated from agar plates. No SNPs, or short insertions/deletions were identified in mutants 2, 3, 10, 12, 13, and 19.

| Mutant | Compound 18 MIC (μM) | Mutation |
| --- | --- | --- |
| Wild-type | 25 | - |
| 1 | >200 | *rny* P280L |
| 2 | 200 | - |
| 3 | >200 | - |
| 4 | >200 | *rny* P280L |
| 5 | >200 | *rny* P280L |
| 6 | >200 | *rny* P280L |
| 7 | >200 | *rny* P280L |
| 8 | >200 | *rny* P280L |
| 9 | >200 | *rpsJ* A122V |
| 10 | >200 | - |
| 11 | 200 | *rny* P280L |
| 12 | >200 | - |
| 13 | 200 | - |
| 14 | >200 | *rny* P280L |
| 15 | >200 | *rny* P280L |
| 16 | 50 | *rny* G240D, LNEJMEBC_01294 I21N |
| 19 | 100 | - |

(12) Marco, A. B.; Martínez de Baroja, N.; Franco, S.; Garín, J.; Orduna, J.; Villacampa, B.; Revuelto, A.; Andreu, R. Dithienopyrrole as a Rigid Alternative to the Bithiophene π Relay in Chromophores with Second-Order Nonlinear Optical

Properties. *Chemistry An Asian Journal* **2015**, *10* (1), 188–197. https://doi.org/10.1002/asia.201402870.
(13) Vanallan, J. A.; Reynolds, G. A.; Petropoulos, C. C.; Maier, D. P. Reactions of Some 4-methylene-4 *H* -pyran Derivatives with Primary and Secondary Amines. *Journal of Heterocyclic Chem* **1970**, *7* (3), 495–507. https://doi.org/10.1002/jhet.5570070306.
(14) Tejeda-Orusco, V.; Blais, M.; Cabanetos, C.; Blanchard, P.; Andreu, R.; Franco, S.; Orduna, J.; Diosdado, B. E. 4H-Pyranylidene-Based Small Push-Pull Chromophores: Synthesis, Structure, Electronic Properties and Photovoltaic Evaluation. *Dyes and Pigments* **2020**, *178*, 108357. https://doi.org/10.1016/j.dyepig.2020.108357.
(15) Pérez Tejada, R.; Pellejà, L.; Palomares, E.; Franco, S.; Orduna, J.; Garín, J.; Andreu, R. Novel 4*H*-Pyranylidene Organic Dyes for Dye-Sensitized Solar Cells: Effect of Different Heteroaromatic Rings on the Photovoltaic Properties. *Organic Electronics* **2014**, *15* (11), 3237–3250. https://doi.org/10.1016/j.orgel.2014.09.003.
